# Spatial transcriptomics elucidates the central role of cerebral vasculatures in the progression of Japanese encephalitis

**DOI:** 10.1101/2025.03.06.637776

**Authors:** Zhihua Ou, Zhaoyang Wang, Qi Chen, Peidi Ren, Xiuju He, Yan Liang, Ying’an Liang, Jiaxuan Wang, Sha Liao, Dexin Wang, Jie Zhao, Oujia Zhang, Zhenyu Peng, Jianxin Su, Wangsheng Li, Guohai Hu, Ao Chen, Ziqing Deng, Xin Jin, Xun Xu, Junhua Li, Gong Cheng

## Abstract

The cellular and molecular mechanisms underlying the neuropathology of Japanese encephalitis virus (JEV) remain obscure. Herein, we designed Stereo-seq chips to simultaneously capture the *in situ* transcriptomes of both the host and JEV, constructing a comprehensive spatiotemporal pathological landscape for Japanese encephalitis (JE). This study reveals the central role of the vascular system in JE pathogenesis, particularly the meninges, which displayed the strongest signal of inflammation and cell death in the JEV-infected brain. The activation of the *Ackr1*^+^ endothelial cells, disruption of the blood-brain barrier, the migration of JEV-infected monocytes, secretion of immune factors by the infected cells, and the occurrence of pyroptosis and necroptosis form a positive feedback loop, resulting in an increasing influx of immune cells and tissue damage. JEV infection leads to neurological impairments, which may be attributed to the interaction between viral proteins and host cellular factors such as *Stat1*, *Stat3*, *Nfkb1, and Sp1*. As the vascular system serves as a central receiver and amplifier of inflammatory signals, regulating inflammation within the vascular system is essential in mitigating JE progression.

## Introduction

Japanese encephalitis (JE) is an acute inflammatory central nervous system (CNS) disease caused by Japanese encephalitis virus (JEV), which is primarily transmitted by the *Culex* mosquitos^1^. Local transmission of JEV occurs in 24 countries in East Asia and the Western Pacific, putting over 3 billion people at risk of infection^2,3^. Without timely treatment, the condition progresses to encephalitis, characterized by severe fever, neck stiffness, and seizures. Children under the age of 14 account for 75% of the JE patients^3^. 20-30% of JE patients die, while 30-50% of survivors develop persistent neurological or psychiatric sequelae^4^. Although the JEV vaccines are commercially available to reduce JE incidences^5^, infection risk persists for individuals lacking prior vaccinations^6–9^. With no virus-specific treatment available, JEV imposes a continuous burden on public health.

JEV can infect many brain parenchymal regions such as thalamus, brainstem, and hippocampus^10,11^. The replication of JEV in neurons can lead to direct neuronal death, while activation of microglia/glial cells and infiltrating inflammatory monocytes lead to indirect killing of neurons via pro-inflammatory cytokines such as IL-6 and TNF-α^12,13^. Inflammatory Ly6c^hi^ monocytes have paradoxical roles in viral encephalitis. On one hand, monocytes could differentiate into effector cells and produce reactive oxygen species (ROS) which repress and clear the virus. On the other hand, unbalanced and uncontrolled migration and infiltration of monocytes will lead to tissue damage and destruction^14^. However, there exists a significant gap in the comprehensive understanding of the intricate interaction networks among various cell types within the brain during the *in situ* immune response following JEV infection. Moreover, the precise molecular mechanisms modulating the progression of viral infection and the extent of tissue damage remain largely elusive, highlighting the need for further investigation and elucidation.

Spatial transcriptomics (ST) technology can capture transcripts *in situ*, which can facilitate the identification of brain regions and cell types infected by JEV, revealing cell-cell interactions involved in JE pathogenesis. As a member of the *Flavivirus* genus, JEV characteristically lacks a polyA tail, posing a detection hurdle for mRNA capture techniques that predominantly utilize polyT probes. In this study, we design Stereo-seq chips with JEV probes that can simultaneously detect JEV and host transcriptomes to dissect the spatial and temporal transcriptional features of neurological lesions during JEV acute infection phase, identify the key cells, pathways, and inflammatory factors associated with brain damage, providing clues on the diagnosis and treatment of JE.

## Results

### Tailor-made Stereo-seq chips reveal JEV infection profiles in the mouse brain

The original Stereo-seq technology utilizes polyT probes to capture mRNA^15^. However, the mRNA of JEV lacks a polyA tail, which cannot be readily detected. To facilitate the capture of JEV transcripts, we strategically designed a virus-specific probe (VSP) targeting a conserved region in the NS5 gene (5’->3’: AACATGATGGGAAAAAGAGA) of multiple flaviviruses (including JEV, Zika virus, and Dengue virus). The VSPs were integrated into the existing polyT probes at a ratio of 1:10 on the Stereo-seq chips (Fig. 1a), thereby enabling simultaneous capture of both host and viral transcripts. Data from Stereo-seq experiments using JEV-infected mouse brain tissues showed that the captured reads were mainly attributable to polyT and VSPs, with polyT probes capturing approximately 60-80% of the overall reads, mostly mouse transcripts (Extended Data Fig. 1a). When it comes to the detection of JEV transcripts, the VSPs captured over 80% of the reads (Extended Data Fig. 1a). Moreover, the VSPs predominantly captured the *NS5* region, which is in line with our probe design targets, while polyT preferentially captured the *Capsid (C)* region, with *NS5* being the secondary target (Extended Data Fig. 1b). Overall, the VSP was effective in capturing JEV transcripts, enabling viral detection in tissues.

**Fig. 1.**
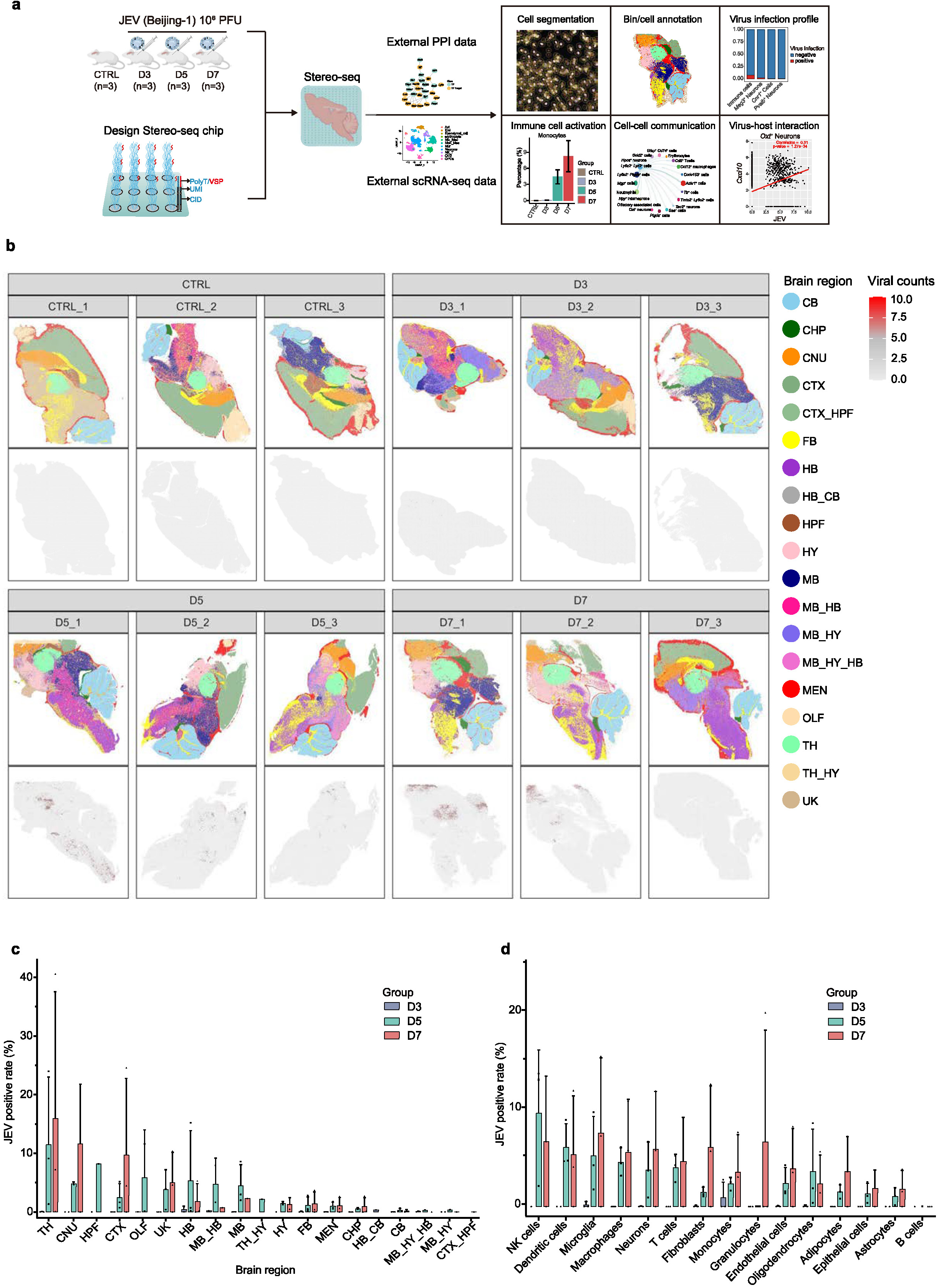
Stereo-seq reveals JEV infection profiles in the mouse model. **a,** Experimental design. 4-week-old female BALB/c mice were intraperitoneally (i.p.) injected with 10^6^ PFU of JEV in 200Cμl of phosphate-buffered saline (PBS) or 200Cμl of PBS in mock-treated mice. Samples were collected at 4 timepoints: control, 3 days post infection (dpi), 5 dpi, and 7 dpi. n indicates the number of samples. A virus-specific probe (VSP) was designed and loaded on the Stereo-seq chips to capture JEV mRNA. The distribution ratio of polyT and VSP probes was 10:1. Mouse brain tissues were collected freshly and embedded in OCT blocks and subjected to H&E staining and Stereo-seq experiment. Besides the Stereo-seq data, external single-cell RNA sequencing (scRNA-seq) data and protein-protein interaction (PPI) data were also included for downstream analysis, including cell segmentation, bin/cell annotation, virus infection profile, cell proportion variation, cell-cell communication, and virus-host interaction. **b,** Spatial distribution of JEV reads and brain region annotation in each sample (Stereo-seq bin35 data, n=12). TH, Thalamus; CTX, Cerebral cortex; CNU, cerebral nuclei; HB, hindbrain; FB, fiber tracts; CHP, choroid plexus; MB, midbrain; HY, hypothalamus; CB, cerebellum; HPF, hippocampal formation; MEN, meninges; OLF, olfactory bulb; UK, unknown. **c,** JEV positive proportions in different brain regions (the number of JEV positive bins divided by the total number of bins in the corresponding brain region) on 3 dpi (D3), 5 dpi (D5), and 7 dpi (D7). The shapes of the dots for D3, D5, and D7 are solid circles, squares, and triangles respectively, and each dot represents one corresponding brain region in that group (Stereo-seq bin35 data). TH, Thalamus; CTX, Cerebral cortex; CNU, cerebral nuclei; HB, hindbrain; FB, fiber tracts; CHP, choroid plexus; MB, midbrain; HY, hypothalamus; CB, cerebellum; HPF, hippocampal formation; MEN, meninges; OLF, olfactory bulb; UK, unknown. **d,** The JEV positive rate for different cell types on D3, D5, and D7 (Stereo-seq bin35 data, n=3 for each group).

To explore JEV infection dynamics in the mouse brain, we inoculated BALB/c mice with 10^6^ PFU of JEV in 200 μl of phosphate-buffered saline (PBS) or 200 μl of PBS in the mock-treated group (Fig. 1a). On the 3^rd^, 5^th^, and 7^th^ day post inoculation (dpi), the mouse brains (n = 3) were collected for Stereo-seq experiment. Based on the overall data quality, we chose an analytical unit of bin35, representing a square region of 35 spots x 35 spots, equal to 17.22 μm x 17.22 μm. The median number of genes detected for each bin ranged from 396 to 844, with an average of 671 genes (Supplementary Table 1). The spatial transcriptomics data showed that the JEV genes were expressed in approximately 0.04%, 3.49%, and 4.25% area of the brain tissues on 3, 5, and 7 dpi respectively, reaching a peak on 7 dpi (Supplementary Table 1). To determine the specific regions infected by JEV, we annotated the mouse brain regions using the marker genes and anatomical information from the Allen Brain Atlas (https://atlas.brain-map.org/, Supplementary Table 2). For instance, the thalamus was annotated using marker genes such as *Prkcd*, *Tcf7l2*, and *Ntng1*, while the meninges were annotated by genes such as *Tmem212*, *Igfbp4*, *Ccdc153*, and *Ptgds.* Our data showed that the thalamus, cerebral cortex, and cerebral nuclei were the most infected regions by JEV (Fig. 1b,c). On 3 dpi, most brain regions were negative for JEV infection. On 5 dpi, multiple brain regions were JEV positive, but the percentages of virus positive bins in thalamus, cerebral cortex, and cerebral nuclei were less than 20%. On 7 dpi, the JEV positive rates in these brain regions rose to 20%-40%, indicating continuous amplification of JEV in these sites.

We further employed SingleR^16^ and its accompanying reference dataset to annotate the cell types and characterized the cellular infection profiles (Extended Data Fig. 1c, Supplementary Table 3). On 3 dpi, monocytes were the dominant JEV positive cells, with an average positive rate of 0.93% (Fig. 1d). On 5 dpi, the highest JEV positive rate was observed in NK cells, 9.65%, followed by dendritic cells (6.12%), microglia (5.25%), macrophages (4.60%), and neurons (3.79%). On 7 dpi, the highest JEV positive rate was observed in microglia (7.60%), followed by NK cells (6.73%). The virus positive rates of NK cells and dendritic cells reached an early peak on 5 dpi, while those of the other cell types (microglia, macrophages, neurons, T cells, fibroblasts, monocytes, granulocytes, endothelial cells, oligodendrocytes, adipocytes, epithelial cells, and astrocytes) peaked on 7 dpi. This evidence suggests that monocytes may be the initial infected cells in the CNS and further spread the virus to the resident cell types in the brain. The phagocytosis of JEV-infected cells and natural viral infection may collectively contribute to the high positive rate of multiple immune cells.

### Spatial transcriptomics reveals the immunological landscape during JE progression

To gain a comprehensive view of the cerebral immunological and physiological alterations following JEV infection, we conducted a comparative analysis to identify the temporal transcriptional changes. A total of 225 genes were found to be differentially expressed between the infection and control groups (Supplementary Table 4). Chronologically, 35, 203, and 112 genes were upregulated on 3, 5, and 7 dpi, respectively (Extended Data Fig. 2a), with 73 genes upregulated on both 5 dpi and 7 dpi. Meanwhile, a total of 58 genes were downregulated after infection, with 2, 6, and 56 genes downregulated on 3, 5, and 7 dpi respectively (Extended Data Fig. 2b). Gene Ontology (GO) enrichment analysis revealed that the upregulated genes in JEV infection groups were mainly associated with immune response (Fig. 2a), while the downregulated genes were mostly associated with functional pathways of brain cells including exocytosis, synapse organization, and neurotransmitter transport, indicating functional damage in the brain (Fig. 2b). Starting from 3 dpi, innate immune response was initiated to recruit lymphocytes, produce interferons and cytokines, induce cell killing, and inhibit virus replication. We observed an early activation of type I interferons (IFN-α and IFN-β) starting from 3 dpi, followed by the activation of type II interferon (IFN-γ) on 5 dpi (Fig. 2a). Stimulated by virus infection, expression levels of multiple Toll-like receptors were enhanced since 3 dpi (Extended Data Fig. 2c), especially *Tlr8*, *Tlr2*, *Tlr3* and *Tlr7*. A series of interferon regulatory factors (IRFs) were also upregulated (Extended Data Fig. 2d), such as *Irf7*, *Irf1*, *Irf8*, *Irf9*, *Irf5*, and *Irf4*. The expression of these genes mostly peaked on 5 dpi. Consistently, the interferon genes were initially expressed on 3 dpi, peaked on 5 dpi, and were downregulated on 7 dpi (Extended Data Fig. 2e). Consequently, the interferon-inducible genes showed low expression levels on 3 dpi but higher levels on 5 dpi and 7 dpi (Extended Data Fig. 2f). On 5 dpi, the pathways including Toll-like receptor signaling, ribosome assembly, protein folding, and cytoplasmic translation were uniquely upregulated (Fig. 2a), which may be linked to the active replication of JEV in the brain. More immune cell types were activated on 5 dpi, including astrocyte development, glial cell differentiation and migration, chemotaxis of granulocytes and lymphocytes (Fig. 2a). On 7 dpi, the immune response was further strengthened by B cell mediated immunity and the activation of myeloid leukocytes and macrophages (Fig. 2a).

**Fig. 2.**
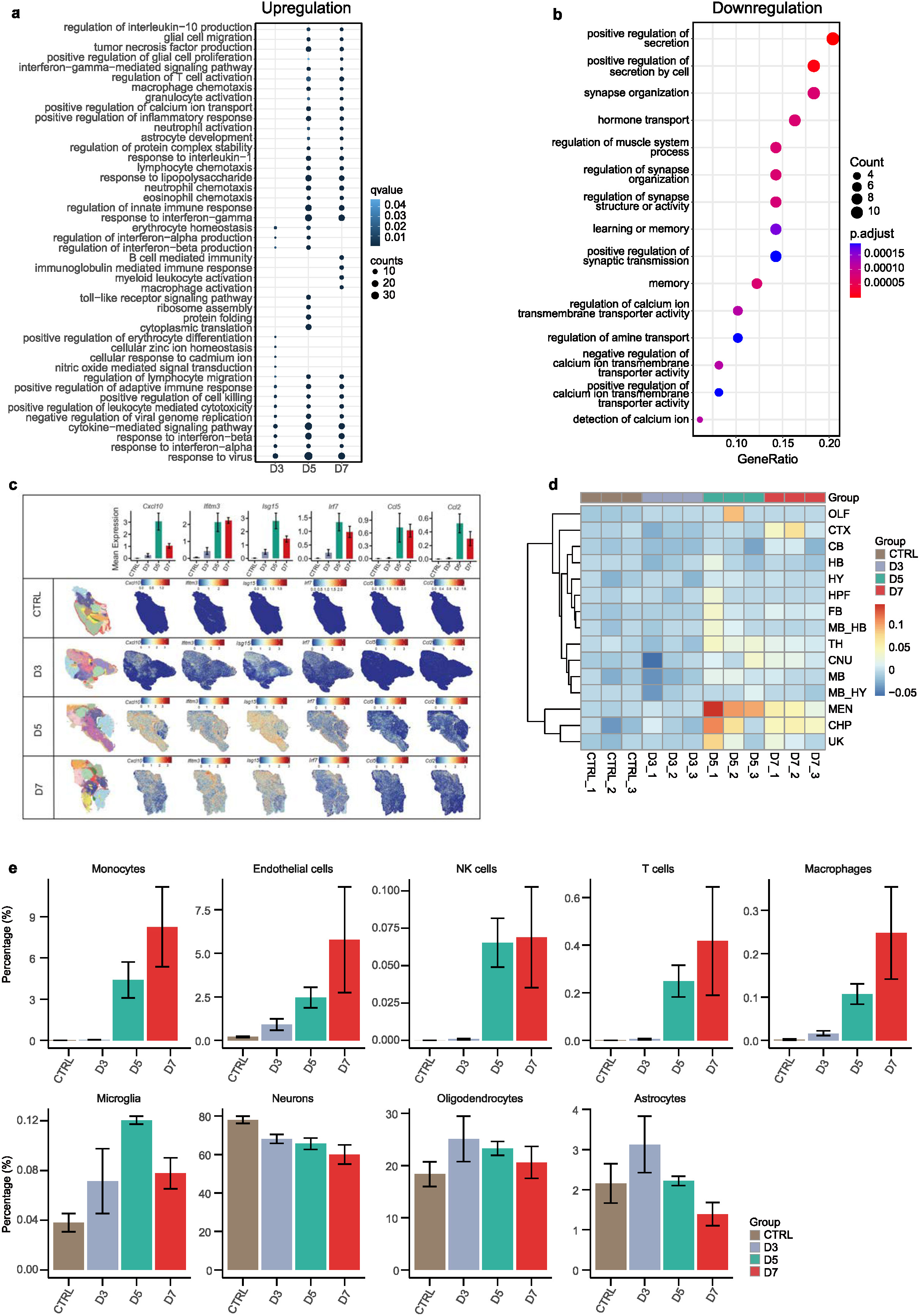
Transcriptional and cellular changes induced by JEV infection in the mouse brain. **a,** Pathway enrichment of upregulated genes on 3, 5, and 7 dpi compared to the control group. Data based on the entire chip (Stereo-seq bin35 data, n=3 for each group). **b,** Pathway enrichment of downregulated genes in the infection group compared to the control group. Data based on the entire chip (Stereo-seq bin35 data, n=3 for control and n=9 for the infection group). **c,** Spatiotemporal expression of immune genes. Top panel: bar plot showing the expression level of immune gene in different groups (Stereo-seq bin35 data, n=3 for each group), and the values in the bar plot represent the average gene expression levels across each sample. Bottom panel: spatial feature plot showing the spatial expression of inflammatory genes in a representative chip of each group (Stereo-seq bin35 data, Sample names: CTRL_3, D3_1, D5_1, and D7_1. **d,** Heatmap showing the inflammatory scores for each brain region of the 12 samples (Stereo-seq bin35 data). **e,** Comparison of the cell type proportion in the control and infection groups including monocytes, endothelial cells, NK cells, T cells, macrophages, microglia, neurons, oligodendrocytes, and astrocytes (Stereo-seq bin35 data, n=3 for each group).

Spatial mapping of the upregulated immune genes such as *Cxcl10*, *Ifitm3*, *Isg15*, *Irf7*, *Ccl2*, and *Ccl5* revealed significantly high expression levels along the vasculatures in the brain (Fig. 2c). To understand the immune response status of different brain regions, we conducted Gene Set Variation Analysis (GSVA) for each brain sample based on the pro-inflammatory gene set from the GSEA database^17–19^ (Supplementary Table 5). Our data showed that the meninges and choroid plexus had the highest immune response scores across different infection timepoints, followed by the brain regions with high infection rates such as the thalamus and cerebral cortex (Fig. 2d), especially on 5 and 7 dpi. At cellular level, comparative temporal analysis of whole Stereo-seq chips demonstrated an increase in the numbers of monocytes, macrophages, T cells, and microglia, in contrast to a progressive decline in the number of neurons (Fig. 2e). We also observed increased signals of endothelial cells (Fig. 2e), which was probably due to vascular dilation caused by immune infiltration or tissue edema, instead of cell proliferation. These results provide a comprehensive overview of the gene expression and biological changes in the JEV-infected brain.

### *Ackr1^+^* endothelial cells actively recruit immune cells and accelerate inflammation in the brain after JEV infection

Regarding the intense involvement of the meninges in JE progression (Fig. 2c,d), we extracted the spatial transcriptomics data of meninges from all samples for in-depth investigation. Analysis of differentially expressed genes (DEGs) showed that many inflammatory molecules including *Cxcl10* and *Ccl2*, as well as MHCII-related genes (*B2m* and *H2-K1*) were upregulated in samples collected on 5 and 7 dpi, except that the upregulation levels on 5 dpi were higher than 7 dpi (Extended Data Fig. 3a). We further conducted cell clustering and annotations using the bin35 data of meninges (Extended Data Fig. 3b,c). By sub-clustering the *Ly6c2*^+^ cells, we identified a cluster of brain endothelial cells (ECs) highly expressing *Ackr1*, defined as *Ackr1^+^* ECs (Fig. 3a). The *Ackr1^+^* ECs highly expressed marker genes that were enriched in angiogenesis and vascular pathways (Extended Data Fig. 3d), including *Vwf*, *Igfbp7,* and *Ctla2a*^20^ (Fig. 3b). Sequential comparative analysis showed an increase in the proportion of *Ackr1^+^*ECs after infection, especially on 5 and 7 dpi, along with significant infiltration of *Ly6c2*^+^*Lyz2*^+^ monocytes and *CD8^+^* T cells in the meninges on 5 dpi (Fig. 3c). Cell-cell communication^21^ analysis of the meninges revealed strong interactions between *Ackr1*^+^ ECs and *Ly6c2*^+^*Lyz2*^+^ monocytes via the CCL/CXCL pathway (Fig. 3d,e,f), which was significantly upregulated on 5 dpi and 7 dpi (Fig. 3d). The *Ackr1*^+^ ECs may recruit *Ly6c2*^+^*Lyz2*^+^ monocytes highly expressing *Cxcl9*, *Cxcl10*, *Ccl7*, *Ccl*5 and *Ccl2* (Fig. 3g). Moreover, the *Ly6c2*^+^*Lyz*2^+^ monocytes also expressed a high level of *Ackr1, Ccr2*, *Ccr1*, and *Ccr5*, which may further facilitate the recruitment of monocytes and T cells (Fig. 3g). Spatially, we observed strong co-localization signals for ligand-receptor pairs including *Ccl7*-*Ccr2*, *Ccl2*-*Ccr2*, *Ccl5*-*Ccr1*, *Ccl2*-*Ccr5 and Cxcl9*/*Cxcl10*/*Ccl7*/*Ccl*5/*Ccl2-Ackr1* in the meningeal region on 5 and 7 dpi (Fig. 3h). Regulatory factor analysis^22^ with the cytokines expressed by *Ly6c2*^+^*Lyz2*^+^ monocytes revealed that the downstream target genes in *Ackr1*^+^ ECs included *Irf7, Ifit3*, *Ccl12*, *Ccl4*, and *Ccl7*, which can further promote the production of proinflammatory cytokines and chemokines by *Ackr1*^+^ ECs (Fig. 3i,j).

**Fig. 3.**
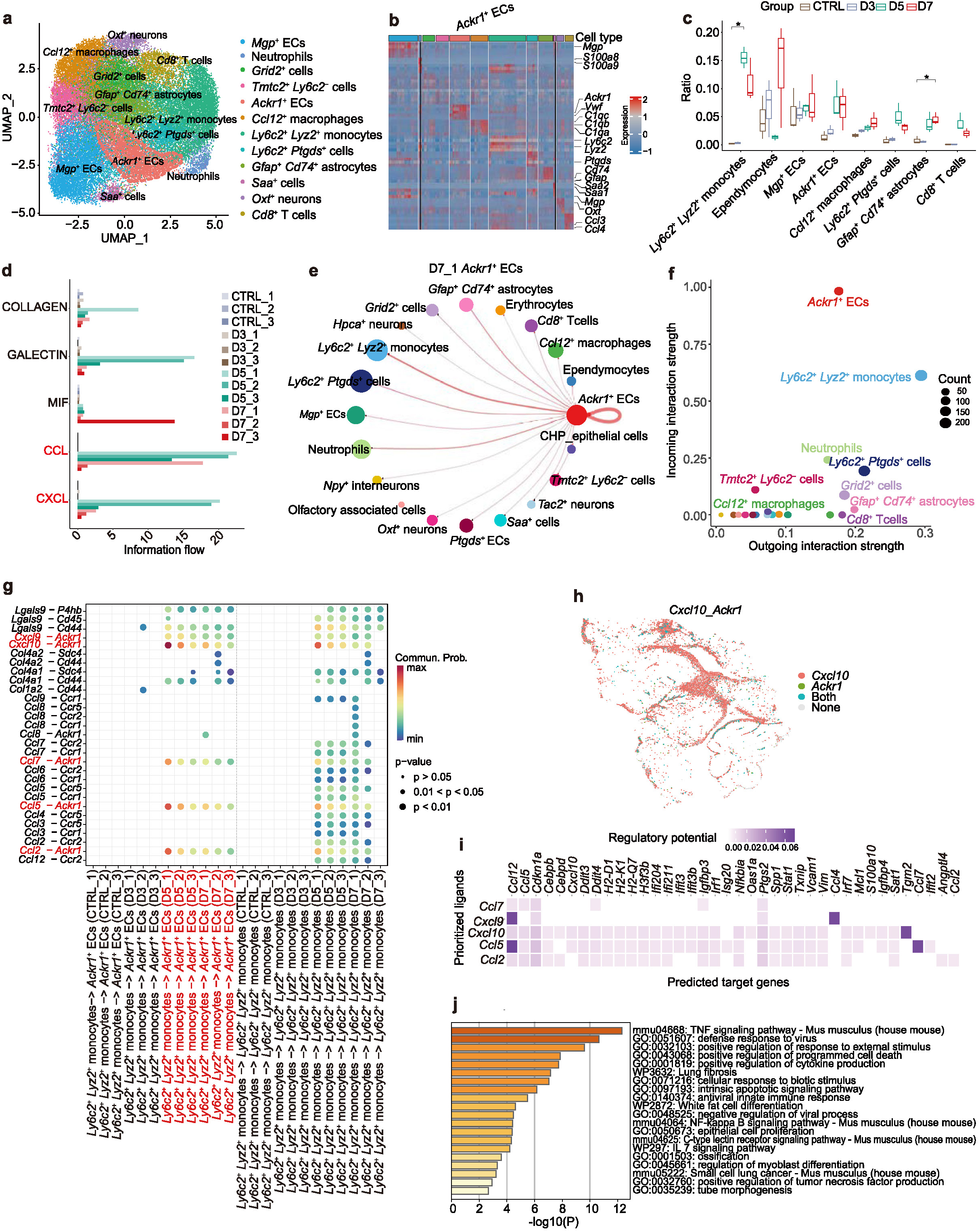
Cell composition and cell-cell interactions in the meninges of JEV-infected mice. **a,** UMAP showing the subpopulations of *Ly6c2*^+^ cells in mouse meninges (Stereo-seq bin35 data, n=12). **b,** Heatmap showing the marker genes of each subpopulation of *Ly6c2*^+^ cells. The cell types share the same color code with (a). **c,** The ratio of different cell types in CTRL (control), D3 (3 dpi), D5 (5 dpi), and D7 (7 dpi) groups (Stereo-seq bin35 data, n=3 for each group). **d,** Intensity flow of each signaling pathway involved in the cell-cell interactions identified in different samples (Stereo-seq bin35 data, n=3 for each group). **e,** The cell-cell interactions between *Ackr1*^+^ cells and the other cell types in sample D7_1 (Stereo-seq bin35 data). **f,** Cell-cell communication strength in the CCL pathway inferred by CellChat in sample D5_2 (Stereo-seq bin35 data), which shows that *Ly6c2*^+^ *Lyz2*^+^ monocytes mainly release the signal while *Ackr1*^+^ cells receive the signal. **g,** Bubble chart showing the ligand-receptor pairs between *Ly6c2*^+^ *Lyz2*^+^ monocytes and *Ackr1*^+^ cells (Stereo-seq bin35 data, n=3 for each group). **h,** The spatial distribution of *Cxcl10*-*Ackr1* ligand-receptor pair in sample D7_1 (Stereo-seq bin35 data). **i,** Heatmap showing the targets in *Ackr1*^+^ cells that potentially regulated by the ligands of *Ly6c2*^+^ *Lyz2*^+^ monocytes (Stereo-seq bin35 data, total n=12). **j,** Bar plot showing the enriched pathway in the *Ackr1*^+^ cells after interacting with *Ly6c2*^+^ *Lyz2*^+^ monocytes (Stereo-seq bin35 data, n=12).

Disruption of the blood-brain barrier (BBB) is crucial for immune cell infiltration and virus invasion into the brain. On 5 and 7 dpi, the tight junction associated genes such as *Tjp1* and *Cldn5* were downregulated (Extended Data Fig. 3e), suggesting the impairment of the BBB. Moreover, the chemokine receptor genes (*Ackr1, Ccr2,* and *Ccr5)* as well as adhesion molecule genes (*Icam1* and *Vcam1)* were also upregulated on 5 and 7 dpi in the meninges, which may promote the transendothelial migration of monocytes and other immune cells^14^ (Extended Data Fig. 3e). Notably, besides a small number of viral RNA detected in the ECs (Fig. 1d), we also observed the upregulation of a series of pattern recognition receptors^23–25^ (PRRs) in different types of ECs, including *Tlr1, Tlr2, Tlr3, Tlr4, Tlr6, Tlr7, Tlr13, Ifih1,* and *Ddx58*, suggesting potential JEV infection in the ECs (Extended Data Fig. 3f). The ECs also displayed higher interferon production activities, as the gene expressions of interferon regulatory factors and interferon receptors were also upregulated in these cells (Extended Data Fig. 3f).

To obtain single-cell resolution evidence, we performed cell segmentation on mouse brains collected from the control (n = 2) and 5 dpi (n = 1) groups and extracted the segmented cells in the meningeal region. Similar to the results based on bin35 data, *Ackr1^+^* ECs were identified in the meningeal region collected on 5 dpi (Extended Data Fig. 4a,b,c). Regulatory analysis with Pyscenic^26^ showed that *Ackr1*^+^ ECs exhibited activation of a series of specific transcription factors (Extended Data Fig. 4d), among which *Irf7*, *Stat1, Cebpb, Cebpd, Spi1,* and *Irf1* were highly expressed in the *Ackr1^+^* ECs (Extended Data Fig. 4e), with *Irf7* displaying the highest expression level. Cell-cell communication analysis with the segmented cell data further confirmed the interactions between *Ackr1^+^* ECs and *Ly6c2^+^Lyz2^+^* monocytes through the *Cxcl10*/*Cxcl9*/*Ccl2*/*Ccl5*/*Cc7*-*Acrk1* axes (Extended Data Fig. 4f,g). Remarkably, the single-cell resolution data showed that among the three types of ECs, *Ackr1*^+^ ECs significantly upregulated the expression of genes associated with pathogen recognition, interferon production, and cell adhesion (Extended Data Fig. 4h,i). These results reveal the central role of *Ackr1*^+^ ECs in receiving and amplifying immune signals during the progression of JE.

### Activation and expansion of *Ccl12*^+^ microglia triggered by JEV infection

Microglia account for approximately 5%-20% of the total glia cell population and are vital to pathogen defense in the brain^27^. Indeed, we observed a sharp increase of activated microglia in the mouse brain after JEV infection (Fig. 2e). By examining the expression of pro-inflammatory and anti-inflammatory markers, we observed increased expressions of pro-inflammatory genes including *Cd14, Fcgr3, Fcgr2b, Cd40, Cd86,* and *H2-Ab1* in the microglia, particularly on 5 and 7 dpi (Extended Data Fig. 5a). Pro-inflammatory chemokines such as *Ccl3, Ccl5*, and *Ccl12* were also significantly upregulated in microglia on 5 and 7 dpi (Fig. 4a), suggesting microglia activation. *Ccl12* was specifically expressed in the activated microglia but not in the homeostatic ones (Fig. 4b,c). Therefore, we defined the activated microglia as *Ccl12^+^*microglia. This result was further confirmed by a public single-cell RNA sequencing (scRNA-seq) dataset (GSE237915) ^13^ where *Ccl12^+^* microglia were also identified and showed a strong correlation with those in our spatial transcriptomics dataset (Extended Data Fig. 5b,c,d). The *Ccl12^+^*microglia were mainly identified in the JEV infection groups, with few homeostatic microglia detected on 5 and 7 dpi (Fig. 4d). We further conducted functional enrichment of the DEGs between *Ccl12^+^* microglia and homeostatic microglia. The upregulated genes in *Ccl12^+^* microglia were associated with antiviral response and interferon signaling (Fig. 4e), while the downregulated genes were linked to synapse organization, learning or memory, and cognitive pathway (Fig. 4f). This suggests that *Ccl12^+^* microglia play a role in antiviral responses, but their transcriptional changes are linked to neuropathology. Transcriptional regulation analysis^26^ showed that transcriptional regulators including *Irf7, Stat1, and Cebpd* were active in *Ccl12^+^*microglia (Fig. 4g), suggesting their involvement in microglia differentiation. Spatially, the expression of *Irf7* not only colocalized with *Ccl12^+^* microglia but also overlapped with *Cxcl10* and the meninges (Fig. 4h), indicating a close interplay between immune cell chemotaxis and microglia activation.

**Fig. 4.**
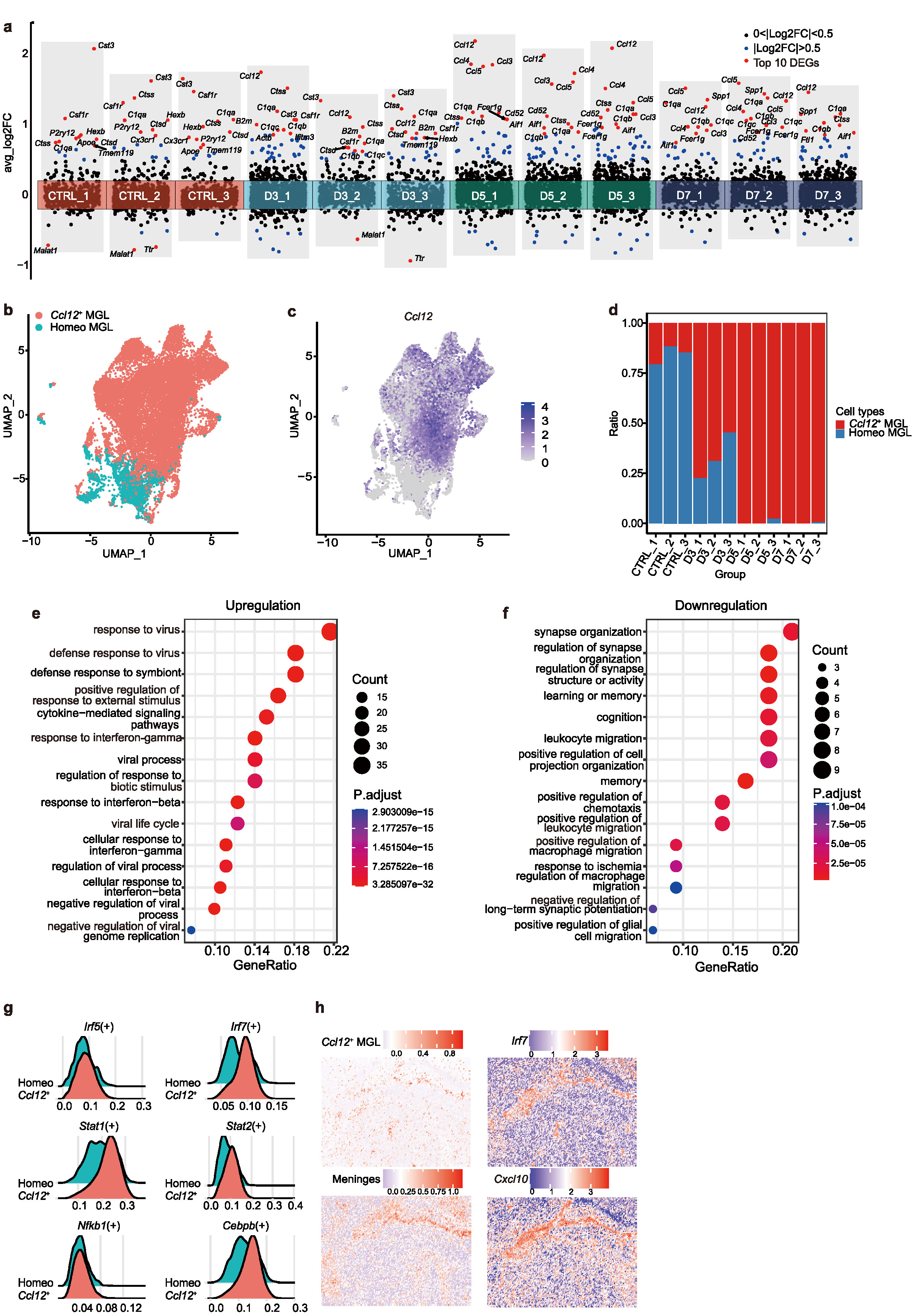
The activation and function of *Ccl12*^+^ microglia in JEV-infected mouse brain. **a,** The differentially expressed genes in microglia compared to all the other cell types of the same sample. **b,** UMAP showing two microglia subtypes based on Stereo-seq bin35 data. The microglia of the 12 samples from the control, D3, D5, and D7 groups were integrated. **c,** UMAP plot showing the expression of *Ccl12* in microglia subtypes (Stereo-seq bin35 data, n=12). **d,** Proportion of homeostatic and *Ccl12*^+^ microglia in each sample (Stereo-seq bin35 data, n=3 for each group). **e-f,** Functional enrichment analysis of the upregulated (e) and downregulated genes (f) in *Ccl12*^+^ microglia compared to the homeostatic microglia (Stereo-seq bin35 data). **g,** Ridge plot showing the upregulation of regulatory factors related to the immune response in homeostatic and *Ccl12*^+^ microglia (Stereo-seq bin35 data, n=3 for each group). **h,** Spatial feature plot showing the signals of *Ccl12*^+^ microglia and meninges, and the expression patterns of *Irf7* and *Cxcl10* in Sample D5_1 (Stereo-seq bin35 data).

### Enhanced necroptosis and pyroptosis in the meninges

To understand the cell death mechanism engaged in JE progression, we estimated the transcriptional score of six death pathways including apoptosis, necroptosis, ferroptosis, cuproptosis, pyroptosis, and autophagy (Supplementary Table 5). Only necroptosis and pyroptosis were found to be associated with JEV infection (Fig. 5a-d, Extended Data Fig. 6a-d). Genes related to necroptosis and pyroptosis were significantly upregulated in brains collected on 5 and 7 dpi (Fig. 5a,c). The signals of necroptosis and pyroptosis were mainly concentrated in the meningeal regions (Fig. 5b,d). Multiple cell types displayed significant necroptosis and signals including *Ackr1^+^* ECs, *CD8^+^*T cells, *Gfap^+^CD74^+^* astrocytes, *Ly6c2*^+^*Lyz*2^+^ monocytes, *Ly6c2*^+^*Ptgds*^+^ cells, neutrophils, and *Saa^+^* cells (Fig. 5e,f), with *Ly6c2*^+^*Lyz*2^+^ monocytes infiltrating from the periphery exhibiting the strongest pyroptosis signals (Fig. 5f). The release of cellular contents due to necroptosis and pyroptosis could enhance the immune cascade reaction, exacerbating neuroinflammation and tissue damage.

**Fig. 5.**
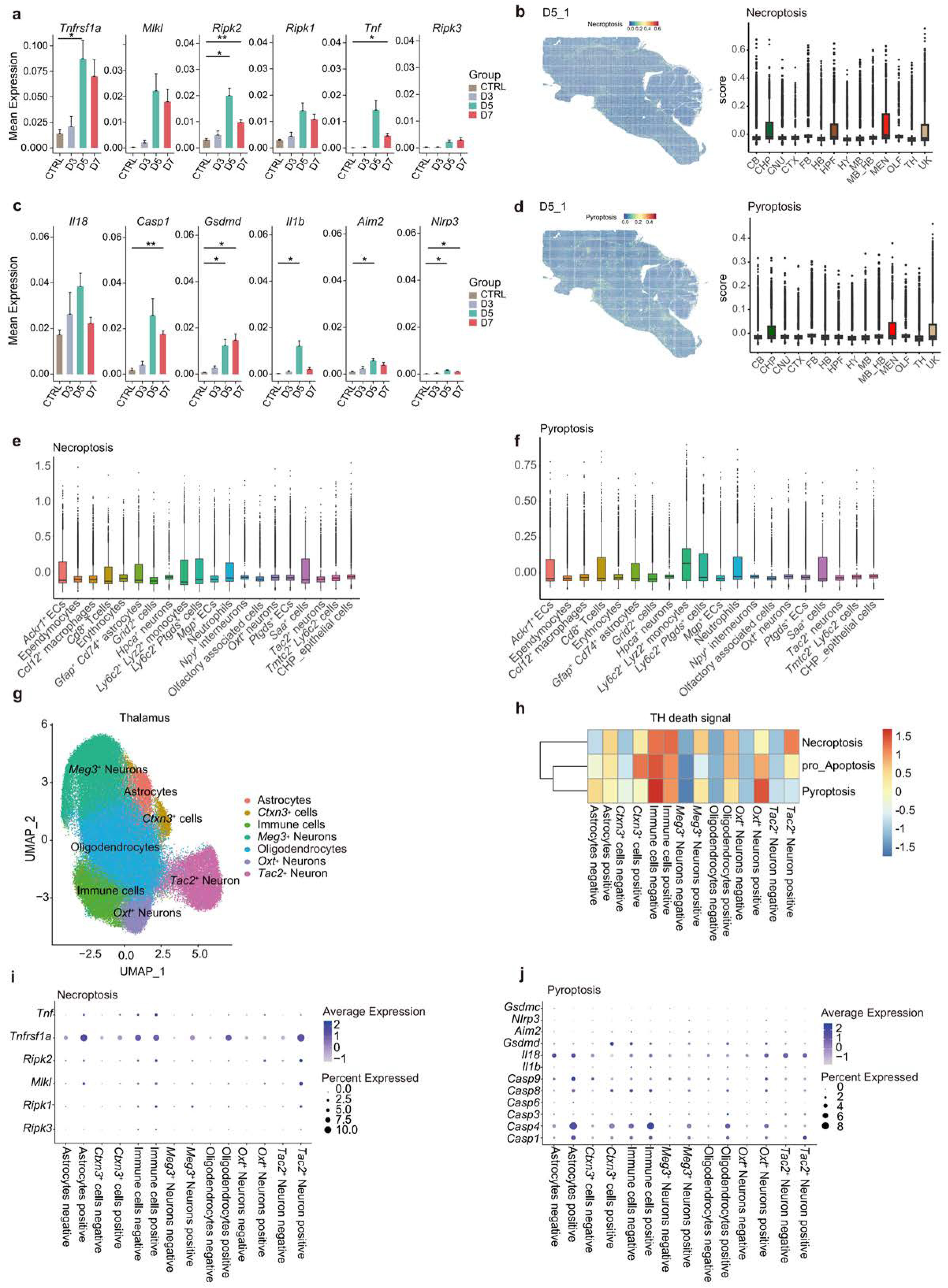
Necroptosis and pyroptosis in the meninges during JEV infection. **a,** Expression levels of necroptosis associated genes in Stereo-seq bin35 data (n=3 for each group). **b,** Spatial intensity of necroptosis gene set signatures in sample D5_1 (left, Stereo-seq bin35 data) and the gene signature scores of necroptosis in different brain regions of D5_1 (right, Stereo-seq bin35 data). **c,** Expression levels of pyroptosis associated genes in Stereo-seq bin35 data (n=3 for each group). **d,** Spatial intensity of pyroptosis gene set signatures in sample D5_1 (left, Stereo-seq bin35 data) and the gene signature scores of pyroptosis in different brain regions of D5_1 (right, Stereo-seq bin35 data). **e-f,** The signature scores of necroptosis (e) and pyroptosis (f) for different cell types in the meninges of JEV-infected samples (Stereo-seq bin35 data). **g,** The UMAP plot of cell types of the thalamus regions from 12 Stereo-seq bin35 samples. **h,** The GSVA scores of necroptosis, pro-apoptosis, and pyroptosis in different cell types and infection states (Stereo-seq bin35 data, n=12). **i-j,** Dot plot showing the expression levels of necroptosis (i) and pyroptosis (j) associated genes in different cell types and infection states (Stereo-seq bin35 data, n=12).

### JEV infection induces necroptosis and pyroptosis in neurons

We further focused on the brain regions with a high viral load such as the thalamus, cerebral nuclei, and hindbrain, to investigate the impact of viral infection on the death of neurons in these regions (Stereo-seq bin35 data). We performed GSVA to evaluate the cell death scores between virus-positive and virus-negative neurons. To avoid potential false positive bins, only bins containing at least two different viral genes or three viral reads were defined as JEV positive. In the thalamus, the JEV-positive *Meg3*^+^ Neurons, *Oxt*^+^ Neurons, and *Tac2*^+^ Neurons exhibited higher pyroptosis and necroptosis signals than their JEV-negative counterparts (Fig. 5g,h). Consistently, genes related to necroptosis (*Ripk3*, *Ripk1*, *Mlkl*, *Ripk2*, *Tnfrsf1a*, and *Tnf*) and pyroptosis (*Casp1*, *Casp4*, *Casp5*, *Casp3*, *Casp6*, *Casp8*, *Casp9*, *Il1b*, *Il18*, *Gsdmd*, *Aim2*, *Nlrp3*, and *Gsdmc*) showed upregulation in the JEV-infected *Oxt*^+^ Neurons and *Tac2*^+^ Neurons (Fig. 5i,j). Similarly, *Pvalb^+^*Neurons in the cerebral cortex (Extended Data Fig. 7a-f), *Oxt^+^*Neurons and *Sst*^+^ Neurons in the cerebral nuclei (Extended Data Fig. 7g-l), as well as *Gpr88*^+^ Neurons and *Sncg*^+^*Nefl*^+^ Neurons in the hindbrain (Extended Data Fig. 7m-r) all upregulated necroptosis and pyroptosis signals in the virus-positive cells. These results suggest that JEV infection causes loss of neurons through pyroptosis and necroptosis (Fig. 5i,j and Extended Data Fig. 7e,f,k,l,q,r). In contrast, the necroptosis and pyroptosis signals in the immune cells were upregulated independent of virus infection (Fig. 5h-j and Extended Data Fig. 7d-f,j-l,p-r), suggesting a more complex cell death mechanism of the infiltrating immune cells.

### Identification of molecules associated with neurological disorders induced by JEV infection

JE patients may experience symptoms such as confusion, aphasia, and paralysis, and the underlying mechanism was not fully explained. To further understand the transcriptional changes related to neurological disorder, we identified modules related to learning, memory, and cognition in multiple brain regions in response to JEV infection using hdWGCNA^28^, including thalamus, cerebral cortex, cerebral nuclei, and hindbrain (Fig. 6a, Extended Data Fig. 8). The functional modules related to learning, memory, and cognition were significantly correlated with specific neuron subtypes, which included the *Meg3*^+^ Neurons in the thalamus (Module SM4, Fig. 6a, Extended Data Fig. 8a-c), the *Meg3*^+^ Neurons in the cerebral cortex (Module SM3, Extended Data Fig. 8d-f), the *Tac1*^+^ Neurons in the cerebral nuclei (Module SM3, Extended Data Fig. 8g-i), and the *Gpr88*^+^ Neurons in the hindbrain (Module SM6, Extended Data Fig. 8j-l). However, the functional genes related to learning, memory, cognition, and locomotion were downregulated in various neuron subtypes rather than restricted to the above-mentioned cell types (Fig. 6b, Extended Data Fig. 9a-c), indicating the extensive neuronal damage caused by JEV infection. For instance, *Ctnx3*^+^ cells, *Meg3*^+^ Neurons, and *Oxt*^+^ Neurons of the thalamus all downregulated genes including *Gnas*, *Cck*, *Snap25,* and *Atp1a3* (Fig. 6b). Comparative analysis revealed that 21 downregulated genes were shared by the learning, memory, and cognition pathways regardless of brain regions (Fig. 6c). Eleven genes of the learning or memory pathway were downregulated in the virus-positive cells in at least three brain regions, with *Atp1a3* and *Ndrg4* identified in six regions (Fig. 6d). Meanwhile, eight genes of the locomotory behaviour pathway were downregulated in the virus-positive neurons in at least three brain regions, with *Atp1a3* and *Kcnd2* identified in six regions (Fig. 6e). In summary, JEV infection affects extensive neuronal functions through disrupting the expression of functional genes in the neurons.

**Fig. 6.**
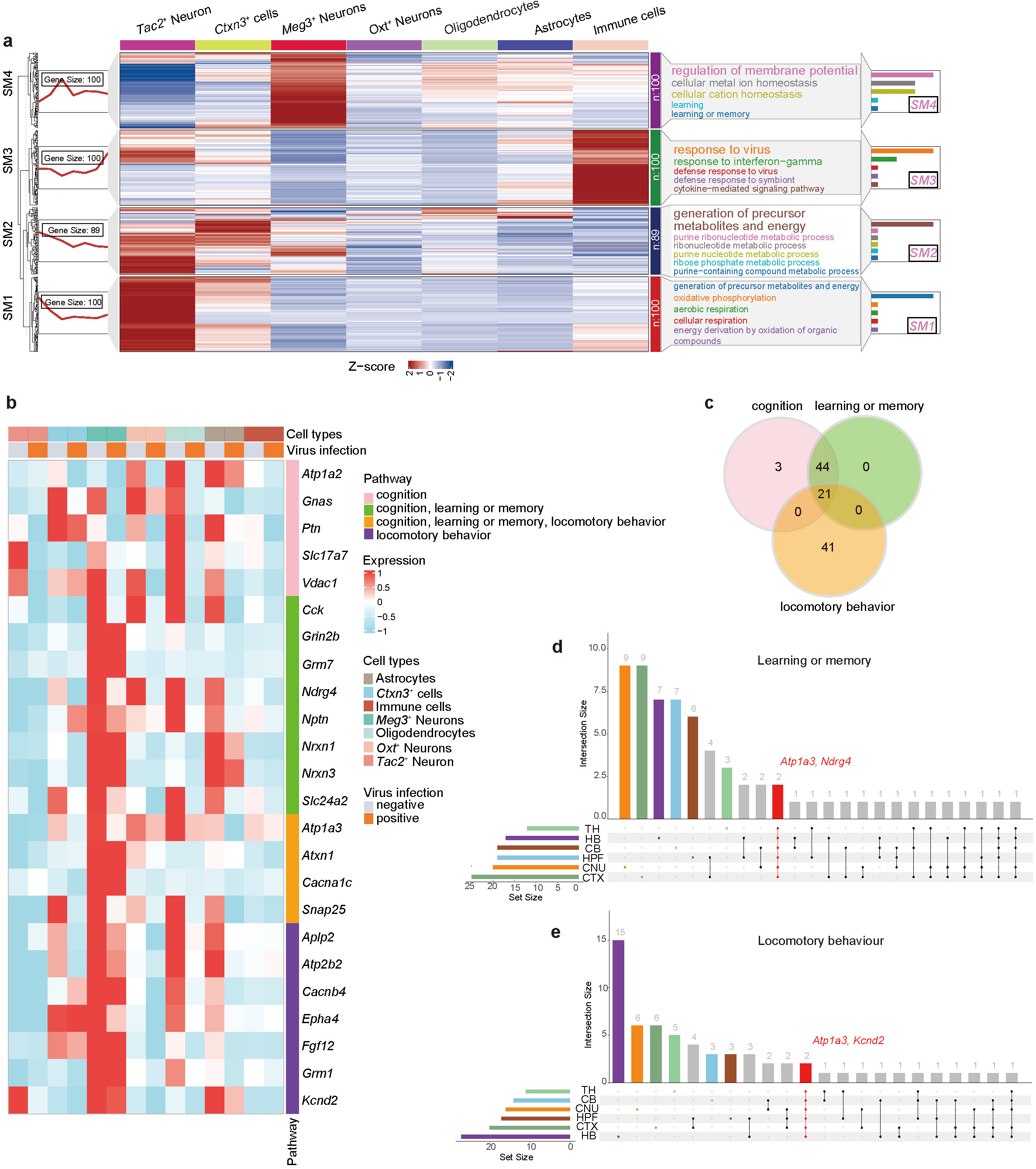
Stereo-seq data reveal the disruption of brain functional modules during JEV infection. **a,** Gene modules identified in the Stereo-seq bin35 data of thalamus. Left panel: Clustering of four major functional modules involved in JE progression. The top 100 genes per module were selected. If the number of genes was smaller than 100, they were all included. Middle panel: Heatmap showing the expression of representative genes in each module for different cell types. Right panel: Representative enriched gene ontology (GO) terms for each module, with each horizontal bar showing the number of genes involved in its corresponding pathway of the same color. **b,** Heatmap showing the expression of genes involved in cognition, learning or memory and locomotory behavior in different cell types and infection states (Stereo-seq bin35 data of thalamus from all samples, n=12). **c,** The Venn diagram showing the overlap of dysregulated functional genes in cognition, learning or memory and locomotory behavior pathway (Stereo-seq bin35 data, total chips=12). Genes were identified based on the comparison between virus-positive and virus-negative cells. The cells were extracted from brain regions with high JEV infection rates including TH, HB, CB, HPF, and CNU. The negative control group was also included for comparison. **d-e,** The upset plot showing the number of disrupted functional genes identified in different brain regions for learning or memory (d), and locomotory behavior (e). Dots with connecting lines indicate overlapped genes between brain regions. *Atp1a3* and *Ndrg4* are involved in learning or memory and both are detected in six brain regions. *Atp1a3* and *Kcnd2* are involved in locomotory behavior and both are detected in six brain regions (Stereo-seq bin35 data of TH, HB, CB, HPF, CNU and CTX, total n=12, 8, 12, 5,10, and 12 respectively).

### Interactions between JEV and host factors

With both virus and host transcriptome detected, we sought to explore JEV and host interactions through integrative analysis. Currently, there exist few systematic studies on the protein-protein interactions (PPIs) between JEV and its host, including experimental or computational prediction. We employed an unsupervised sequence embedding method (doc2vec) and random forest classifier to identify potential PPIs between JEV and human ^29^. A total of 588 human proteins interacting with the virus were predicted (Supplementary Table 6). There were 550 proteins interacting with NS5 and 490 with NS3 (Extended Data Fig. 10a). By performing pathway enrichment analysis on the host proteins interacting with each viral protein, we found that each viral protein uniquely perturbed 2-7 pathways except for C protein whose interacted host proteins always overlapped with those of the other viral proteins (Extended Data Fig. 10b). The NS5 protein affects pathways related to protein stability and cell cycle mitosis, while NS3 specifically hijacks pathways related to endoplasmic reticulum processing of proteins (Extended Data Fig. 10b). The pathways commonly associated with at least seven virus proteins included antiviral response of interferon-stimulated genes, protein localization to organelles, virus infection, and RNA metabolism. Pathway enrichment of the host proteins interacting with all the virus proteins reveal pathways including response to multiple infections, cellular response to interleukin-4, rhythmic process, and HDAC6 interactions in the CNS (Extended Data Fig. 10c, Supplementary Table 7). Among them, the HDAC6 interaction and Parkin ubiquitin proteasome pathway may be involved in nervous system dysfunction. In the Parkin ubiquitin proteasome pathway, members of the HSP70 family, Tubulin family, and 26S ubiquitin proteasome family all extensively interacted with viral proteins.

We further combined PPI results with our spatial transcriptomics data to identify transcriptional changes due to direct virus-host interaction. Because of the low detection level of JEV mRNAs in the spatial transcriptomic data (Supplementary Table 1), the correlation coefficients between host and viral genes were smaller than 0.25 (Extended Data Fig. 11a). Even using a low threshold (|correlation coefficient| >0.2), less than 70 host genes were found to be correlated with virus gene expression in a defined cell type and only 1-3 genes identified in the PPI were included (Extended Data Fig. 11b). When we used the segmented cell data of Sample D5_1, the number of host genes identified in a single cell type correlated with virus replication increased (|correlation coefficient| >0.2) but the overlap with PPI remained minimal (Extended Data Fig. 11c). Similar results were obtained when we conducted correlation analysis (|correlation coefficient| >0.3) using the external scRNA-seq data (GSE237915) ^13^, although the number of overlapped genes increased (Extended Data Fig. 11d,e). Unfortunately, no genes were consistently identified across the three datasets.

While the low capture rate of mRNAs due to technical limitations may mask transcriptomic changes, it’s also possible that the viral proteins may hijack host transcription factors to influence downstream mRNA expression indirectly. For instance, the NS5 protein of Dengue virus can bind the PAF1C complex, thereby affecting the expression of downstream interferon-related genes^30^. Therefore, we intersected the PPI results with known transcription factors, revealing 59 transcription factors that interact with JEV proteins (Extended Data Fig. 11f) along with their downstream target genes. Because the thalamus has the highest virus infection rate among different brain regions, we decided to conduct host-virus interaction analysis using cells of the thalamus. Based on the bin35 data of the thalamus, 18 target genes correlated with viral gene expressions were identified (Extended Data Fig. 11g, (Supplementary Table 8), which were regulated by 13 transcription factors. A total of 158 target genes with expression correlated with viral genes were identified in the external scRNA-seq dataset (GSE237915)^13^, regulated by 20 transcription factors (Extended Data Fig. 11g, Supplementary Table 8). There were 13 transcription factors and 14 target genes shared by these two datasets (Extended Data Fig. 11g), with NS3 and NS5 interacting with nearly all these 13 transcription factors (Fig. 7a). The regulatory network revealed *Nfkb1, Sp1, Stat1*, and *Stat3* as hub transcription factors and *Cdkn1a, Mt1*, and *Cxcl10* as hub target genes (Fig. 7b). The 14 common target genes were enriched in pathways that positively regulate monocyte chemotaxis and spinal cord injury, among which *Cxcl10*, *Gfap*, *Ccl2*, *Mbp*, and *Tspo* are common to both pathways (Fig. 7c). *Cxcl10*, *Gfap*, *Ccl2,* and *Tspo* showed a positive correlation with viral gene expression (Fig. 7d, Extended Data Fig. 12), while *Mbp* displayed a negative correlation (Extended Data Fig. 12). These results reveal key molecules involved in the host response to JEV infection.

**Fig. 7.**
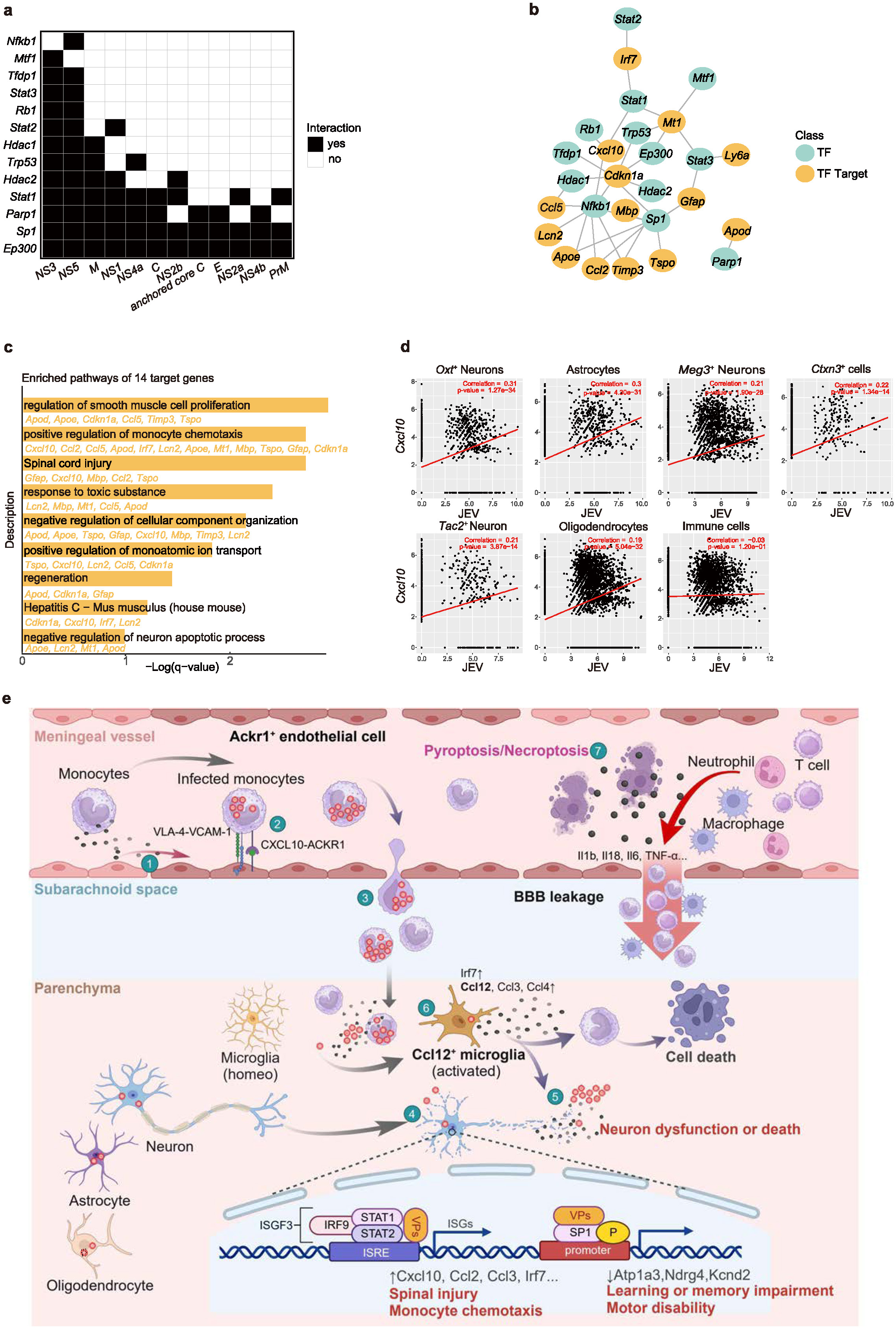
Integrative analysis of PPI and transcriptomic data reveal host gene expression changes affected by JEV infection. **a,** The interaction of 12 virus proteins with 13 transcriptional factors. **b,** Network showing the regulatory relationships of 13 transcriptional factors and 14 corresponding target genes. **c,** Enriched pathways of 14 target genes. **d,** Correlation of JEV transcripts with *Cxcl10* in different cell types, calculated using Stereo-seq bin35 data of thalamus from 9 samples infected by JEV and 3 non-infected samples. **e,** The immunopathological landscape of JE during the acute infection stage. 1) *Ackr1*^+^ endothelial cells in the brain vascular system are activated by the inflammatory factors in blood vessels when the host is infected by JEV. Their tight junctions become damaged and BBB permeability is disrupted. 2) *Ackr1*^+^ endothelial cells help peripheral monocytes infiltrate into brain parenchyma through multiple chemokines-ACKR1 and adhesion related interactions such as VCAM-1-VLA-4. 3) The JEV-infected monocytes disperse the virus into the brain parenchyma through the “Trojan Horse” mode. 4) Multiple cell types in the brain parenchyma are infected by JEV, affecting anatomical regions including the thalamus, cerebral cortex, cerebral nuclei, hindbrain, etc. The infected neurons overexpress pro-inflammatory cytokines (Cxcl10, Ccl2, Ccl3, Irf7) and down-regulate molecules (Atp1a3, Ndrg4, Kcnd2) related to brain functions including cognition, learning, memory, and locomotory behavior. 5) Cell death occurs in the infected neurons, mostly through the pyroptosis and necroptosis pathways. 6) *Ccl12*^+^ microglia are activated to respond to viral infection by releasing pro-inflammatory cytokines such as Ccl2, Ccl3, and Ccl4. 7) The meninges are the inflammation and cell death hotspot in the JEV-infected brain, mostly contributed by the infiltrating Ly6c^+^ monocytes and their death through the pyroptosis and necroptosis pathways. The inflammatory and cell death signals in the meninges further expedite BBB leakage, exacerbating brain infection, immune cell infiltration, and functional impairment.

### Monocytes and vascular endothelial cells synergize to facilitate JE progression

Our data suggest the role of JEV-infected monocytes in spreading the virus to the CNS and initiating encephalitis (Fig. 7e). Following JEV infection of immune cells in the bloodstream, such as *Ly6c2^+^* monocytes, the infected cells release immune and chemokine factors such as Cxcl10, Cxcl9, Ccl2, and Ccl5. The circulation of these inflammatory factors activates the immune response of the vascular system in the CNS, including the meninges and choroid plexus. As a result, a subset of *Ackr1*^+^ endothelial cells are activated. Concurrently, the *Ackr1*^+^ endothelial cells upregulate cell adhesion-related molecules such as *Icam1* and *Vcam1*, and release additional immune chemokines including *Ccl2*, *Ccl7*, and *Cxcl10*, attracting more immune cells to migrate to the CNS. During this process, monocytes carrying the virus cross the BBB and enter the brain parenchyma in a Trojan horse manner, disseminating the virus to neurons, microglia, and the other cell types. The activation of the *Ackr1*^+^ endothelial cells, secretion of immune factors, disruption of the BBB, and the migration of JEV-infected monocytes form a positive feedback loop, resulting in an increasing influx of immune cells together with viruses into the CNS. Later, the activation of microglia and infiltration of myeloid immune cells help phagocytose and eliminate virus-infected cells. However, the subsequent cell death exacerbates inflammation in the CNS, particularly with the occurrence of pyroptosis and necroptosis in the meninges. JEV infection of the neurons induces neuronal damage and cell death through apoptosis and pyroptosis, leading to a range of symptoms, including cognitive, memory, and behavioral dysfunctions. These may be attributed to the interaction between viral proteins and host factors such as *Stat1*, *Stat3*, *Nfkb1, and Sp1*. During JE development, the vascular system serves as a crucial inflammatory signal receiver and amplifier. The resolution of inflammation within the vascular system is essential in mitigating disease progression.

## Discussion

JEV has a notable neuroinvasive capability, enabling it to breach the BBB and enter the CNS. Despite JEV being the leading cause of viral encephalitis worldwide^31^, the molecular mechanisms through which JEV crosses the BBB and induces encephalitis remain poorly understood. In this study, we employed Stereo-seq technology to conduct a detailed characterization of the spatiotemporal transcriptional dynamics in the mouse brain after JEV infection. We discovered the cellular and molecular mechanisms of key pathogenic processes, including viral infection, BBB disruption, immune cell activation and recruitment, and cell death, thereby providing molecular targets for JE prevention and treatment.

The BBB is composed of brain microvascular endothelial cells (BMECs), tight junctions between endothelial cells, the basement membrane, and astrocytic end-feet. It plays a crucial role in maintaining the homeostasis of the brain’s internal environment. A key step in JEV infection of the brain is to breach the BBB, and the mechanisms are still debated. Some studies suggest that JEV can directly infect BMECs, which subsequently release progeny viruses from the basal side into the brain parenchyma^32^. Other studies propose that early JEV-induced inflammatory factors affect BMECs by downregulating tight junctions, thus increasing BBB permeability and facilitating viral entry into the brain^33,34^. Additionally, the virus can infect immune cells (such as macrophages and monocytes) and cross the BBB in a “Trojan horse” fashion^35^. In our spatial transcriptomics data, the BMECs can be classified into *Ackr1*^+^ and *Ackr1*^-^ subtypes. Compared to *Ackr1*^-^ cells, *Ackr1*^+^ cells exhibit significant activation of inflammatory pathways and reduced expression of tight junction proteins, while showing increased expression of adhesion molecules on their surfaces to recruit virus-infected immune cells (Fig. 3d,g and Extended Data Fig. 4g,h). Although only a small number of virus reads were identified in the BMECs (Fig. 1d), the *Ackr1*^+^ ECs upregulated the PRRs (Extended Data Fig. 4i), indicating potential infection by the virus. This evidence supports that JEV may invade the CNS as free particles and in the Trojan horse form. After JEV breaches the BBB, they infect multiple cell types in the brain, including microglia, neurons, fibroblasts, endothelial cells, oligodendrocytes, adipocytes, epithelial cells, and astrocytes (Fig. 1d), which collectively contribute to encephalitis.

The meninges, which are rich in blood vessels and lymphatic vessels, serve as the boundary of the brain and play a crucial role in the interaction between the CNS and the peripheral immune system. In various infectious or non-infectious brain encephalitis, the immune response in the meninges is often the earliest and most intense compared to the brain parenchyma^36^. Our spatial transcriptomics data showed that the meninges and choroid plexus scored higher than the other brain regions in the inflammatory grading (Fig. 2d), which may be contributed by the infiltration of immune cells such as *Ly6c2*^+^*Lyz2*^+^ monocytes (Fig. 3c). The interplay between the peripheral immune cells and the vascular endothelial cells in the brains (meninges and choroid plexus) is essential for propelling the progression of JE (Fig. 3d-g). Therefore, disruption of the molecular interaction axes that engender endothelial cell activation could potentially impede alterations in vascular permeability, curtail the systemic dissemination of pathogens to various organs, and mitigate hyperinflammation. Wang et al. have shown that Axl is involved in maintaining BBB integrity and antagonist against IL-1α released by the peritoneal macrophages can decrease JE pathogenesis in the context of compromised BBB integrity^37^. Our findings show that *Ly6c2*^+^ monocytes can activate *Ackr1*^+^ ECs through *Cxcl10*/*Cxcl9*/*Ccl2*/*Ccl5*/*Ccl7*-*Ackr1* interactions. The inhibition of these interaction axes may abrogate the pathological alterations of endothelial cells. As no drug targeting ACKR1 is available, interrupting the ligands expressed by the immune cells will be crucial to disease intervention.

Monocytes play a controversial role in the pathogenesis of flavivirus infection. Firstly, they are target cells of flaviviruses, facilitating virus dissemination to various organs^38–40^. Secondly, they can exert pro-inflammatory effects, clearing viruses and dead cells^41,42^. However, the associated excessive infiltration and pyroptosis of immune cells can disrupt normal organ function. Consequently, dampening the excessive recruitment of monocytes and reducing pyroptosis can ameliorate the hyperinflammation triggered by flavivirus infection. In this study, the monocytes accumulated within the vessel through multiple chemokine-ACKR1 axes before entering the parenchyma (Fig. 3c,g), similar to the *Toxoplasma gondii* infection^43^. Kim et al. found that Ly6c^hi^ monocytes can be recruited into CNS by the CCR2-CCL2 axis and that the ablation of CCR2 could restrict Ccr2^+^ monocytes entry and enhance the resistance to JE in a mouse model^44^. The above-mentioned chemokines can be expressed by microglia^45^, astrocytes^46^, and neurons^47^. The monocytes can also be recruited by VLA-4-VCAM-1 and LFA-1-ICAM-1 axes by interacting with endothelial cells to exert transendothelial effects^48^. Targeting the cytokines that are highly expressed by a variety of cells will effectively reduce the excessive recruitment of monocytes, for instance, bindarit^49–51^ has been identified as an inhibitor for *Ccl2*, *Ccl7*, and *Ccl8*. It’s also reported that the Janus kinase inhibitors, tofacitinib and upadacitinib, can help reduce the expression of CXCL10 in patients with drug-induced hypersensitivity syndromes^52^. These may potentially serve as candidate immune modulatory drugs for JE.

Microglia are the resident immune cells of the brain, playing a direct role in neurodegenerative and neurological diseases, which are often accompanied by neuroinflammation, synaptic and neuronal loss, and cognitive decline^53^. During the acute infection stage of JE, the *Ccl12^+^* microglia were activated by transcription factors such as *Irf5, Irf7*, and *Stat1* (Fig. 4g). *Irf5, Irf7*, and *Stat1* were also involved in the activation of *Ackr1^+^* ECs (Extended Data Fig. 4d,e) and their expressions can be triggered by chemokines including *Cxcl10, Ccl2,* and *Ccl5* that were highly expressed in the acute infection phase, especially on 5 dpi (Extended Data Fig. 2f). This underscores the pivotal role of peripheral circulatory chemokines in bolstering the interferon signaling pathway, thereby activating cerebral endothelial and microglial cells and exacerbating immune cascade reaction. The *Ccl12^+^*microglia are essential in the pro-inflammatory response upon JEV infection, whether they are involved in the post-infection neuropathological sequelae remains to be explored.

Previous researches have shown that JEV infection can lead to apoptosis^54,55^, ferroptosis^56^, and pyroptosis^57^. Most of the studies focused on the presence of a single form of death caused by JEV infection in multiple cell lines or mouse models. Except for Wang et al. who systematically assessed the expression of key marker genes for multiple death modalities in peripheral peritoneal macrophages after JEV infection using bulk RNA-seq^57^. Taking advantage of the high detection throughput and wide spatial vision provided by Stereo-seq technology^15^, we have systematically assessed six types of cell death at nearly single-cell resolution. We found that the meninges had stronger pyroptosis and necroptosis signals than the parenchyma. *Ackr1^+^* ECs, *CD8^+^* T cells, *Gfap^+^CD74^+^* cells, *Ly6c2*^+^*Lyz*2^+^ monocytes, *Ly6c2*^+^*Ptgds*^+^ cells, neutrophils, and *Saa^+^* cells are the major cell types contributing to these cell death signals in the meninges. In the brain parenchyma, the virus-positive neurons upregulated apoptosis, pyroptosis, and necroptosis signals, while the immune cells upregulated the cell death signals regardless of virus infection status. Due to the intensive rapture of the cell membrane and the release of pro-inflammatory factors, pyroptosis^58^ and necroptosis can further exacerbate inflammatory response in the meningeal area, intensifying BBB breakage through the bystander effect. Our study indicated that pyroptosis and necroptosis of immune cells in the meninges greatly contribute to neuroinflammation caused by JEV infection. Gasdermins^58^ including GSDME^59^ and GSDMD^60^ are executioners for pyroptosis. Disulfiram has been identified as an inhibitor of pore formation by GSDMD^61^, which may help alleviate excessive inflammation during JE.

Clinically, JE can lead to permanent cognitive or motor impairments yet the molecular changes in the relevant brain regions are not clear. We systematically evaluated the genes associated with such disorders after JEV infection. More dysregulated genes associated with learning or memory were identified in the cerebral cortex and cerebral nuclei, while more locomotory behavior associated genes were found in the hindbrain, although some of these genes are not correlated with virus infection level. Moreover, three genes were consistently downregulated among different brain regions, i.e., *Atp1a3*, *Ndrg4*, and *Kcnd2.* Among these, *Atp1a3* and *Ndrg4* were associated with learning and memory or cognition, while *Atp1a3* and *Kcnd2* were linked with locomotory behavior (Fig. 6d,e). The three genes were also found to be significantly downregulated in Alzheimer’s disease^62–66^. Whether rescuing the functions of these three genes could repair the neurological damage by JEV infection remains to be explored.

To understand how the interactions between the virus and host proteins contribute to JE pathogenesis, we predicted the PPIs between human and JEV, and explored the correlation between PPIs and dysregulated host transcripts. Functional enrichment of the PPIs showed that multiple virus proteins are associated with the Parkin ubiquitin proteasomal system and HDAC6 interactions in the CNS. Activation of the Parkin ubiquitin pathway may be associated with the clearance of virus-infected cells through apoptosis, while HDAC6 was found to be associated with virus infection and inflammation regulation^67–72^. Moreover, inhibition of HDAC6 was found to reduce JEV replication and prevent excess inflammation^73,74^. However, few of the related host genes were differentially expressed between the infected and the mock-infected groups. Given that PARKIN and HDAC6 are involved in protein ubiquitination and deacetylation, the viral and host protein interaction may trigger epigenetic or post-translational modifications of Parkin and HDAC6 instead of affecting gene expression levels. By intersecting the PPI results with single-cell and spatial transcriptomics data, we found that the expression levels of most host genes that interact with the viral proteins were not affected, which may be largely due to the sparsity of the spatial and single cell transcriptomic data. However, the viral proteins may affect the expression of their downstream target genes by binding to transcription factor proteins. We took the intersection of Stereo-seq and scRNA-seq dataset and obtained a total of 13 transcription factors that can bind to viral proteins. 14 target genes downstream of them were identified, which were correlated with viral gene expression and involved in pathways such as monocyte recruitment and spinal cord injury. The integration of PPI and transcriptomics data reveals multiple hub transcription factors and target genes in response to JEV infection (Fig. 7b,c). Among them, *Ccl2, Cxcl10, Ccl5, Lcn2, Ly6a,* and *Irf7* displayed significant overexpression (>100 fold) after JEV infection (Extended Data Fig. 12). These molecules may serve as diagnostic or therapeutic candidates for JEV.

In general, this study utilized high-resolution spatial transcriptomics to decipher the spatiotemporal pathological landscape in JEV-infected mouse brains, elucidating the key interactions between immune cells and brain vascular endothelial cells in the pathogenesis of JE. We also integrated genomic and transcriptomic data to explore the molecular interactions between JEV and the host, offering insights into the biological pathways perturbed by viral infection. These discoveries not only furnish novel biomarkers for the diagnosis and therapeutic intervention of JE but also contribute methodological and theoretical insights for the investigation of pathogenic mechanisms of other flaviviruses.

## Methods

### Cells and virus

C6/36 cells (*Aedes albopictus* lava cells) were grown in RPMI 1640 medium (Cat #11875093, Gibco) supplemented with 10% fetal bovine serum (FBS; Cat# 16000-044, Gibco) and maintained at 28°C. Vero cells (African green monkey kidney cells) were raised in minimum essential medium (MEM; Cat# 11095080, Gibco) supplemented with 5% FBS. JEV Beijing strain-1 (JEV) was propagated in C6/36 cells and titrated on Vero cells by plaque assay as previously reported^75^.

### Mice

Four-week-old female BALB/c mice were obtained from Beijing Vital River Corporation. The mice were bred and maintained in a specific pathogen-free animal facility at Tsinghua University, which is accredited by the Association for Assessment and Accreditation of Laboratory Animal Care International (AAALAC). All animal experiments were approved by the Institutional Animal Care and Use Committee of Tsinghua University and conducted in accordance with their guidelines.

### Animal infection experiment

Four-week-old female BALB/c mice were intraperitoneally (i.p.) injected with 10^6^ PFU of JEV in 200 μl of phosphate-buffered saline (PBS) or 200 μl of PBS in mock-treated mice. On the 3^rd^, 5^th^, and 7^th^ day post inoculation (dpi), the whole brain tissues of JEV-infected and mock-treated mice (negative controls) were collected. Briefly, the mice were anesthetized with Avertin (Cat# HY-B1372, MCE), followed by cardiac perfusion with ice-cold PBS to thoroughly remove the residual blood. After cardiac perfusion, the whole brain tissues were taken out for immediate cryo-embedding with OCT compound (Cat# 4583, SAKURA). Afterward, the brain tissues were stored at -80 ° C for subsequent cryo-section.

### Quality control of OCT-embedded samples

For each OCT-embedded sample, 100-200 μm thick sections were cut and harvested to extract total RNA using the RNeasy Mini Kit (Qiagen, USA) according to the manufacturer’s protocol. The RNA integrity number (RIN) was examined by a 2100 Bioanalyzer (Agilent, USA) and samples with RIN≥7 were selected for downstream experiments. For each sample, one tissue section (10-μm thick) with a large coverage along the anterior-posterior axis containing the hippocampus was preserved for Stereo-seq experiment. One adjacent section from each sample was applied to hematoxylin and eosin (H&E) (Beyotime Biotechnology, China) for pathohistological observation.

### H&E staining

Briefly, the slides were fixed with ice-cold acetone and then rinsed with PBS to remove OCT embedding medium. Afterward, hematoxylin and eosin were used to stain the sections. After staining, the sections were dehydrated in an increasing concentration gradient of ethanol, transparentized with xylene, and sealed with neutral resin.

### Stereo-seq library preparation and sequencing

The spatial transcriptomics experiment was performed using the STOmics gene expression assay kit (BGI-Shenzhen, China). The frozen brain tissue sections were placed onto the Stereo-seq chip surface patterned with DNA nanoballs (DNBs, i.e., spot) and incubated at 37°C for 3 minutes. Then, the sections were fixed in methanol and incubated for 40 minutes. After that, the sections were stained with nucleic acid dye (Thermo Fisher, Q10212) and imaging was performed with a Motic microscope scanner (Motic, China) at the channel of FITC. After washing off the staining solution, the sections were permeabilized for 12 minutes in an incubator at 37°C to allow mRNA *in situ* hybridization. Reverse transcription (RT) was performed at 42°C for 1 hour following mRNA capture. The RT products were subsequently subjected to cDNA release enzyme treatment overnight at 55°C. Then the released cDNA was purified using DNA clean beads and amplified with PCR mix. The PCR products were used for library construction and finally sequenced with a 50 + 100 bp strategy on an MGI DNBSEQ T series sequencer (MGI, China).

### Preliminary processing of Stereo-seq data

Stereo-seq raw data were automatically processed using the BGI Stereomics analytical pipeline (http://stereomap.cngb.org/), where the reads were sequentially processed by barcode demultiplexing, adapter filtering, reference mapping, deduplication, and quantification. The reference genome was a combination of the mouse (*Mus_musculus*, GRCm38) and the JEV (Accession Number: L48961) reference genome. To enhance resolution and ensure that each analytical unit contains more than 200 genes, we merged adjacent DNBs to one bin35 (35 x 35 DNBs, i.e., 17.22 x 17.22 μm). For quality control, the bins with UMIs fewer than 100 and those genes expressed in fewer than three cells were removed. Then the data were normalized, logarithmically transformed, and scaled by Scanpy^76^. Dimension reduction was performed using PCA. Unsupervised clustering of bins was performed using leiden with default parameters in Scanpy^76^. Data statistics are shown in Supplementary Table 1.

### Cell segmentation

Cell segmentation analysis was performed using StereoCell (V1.2.6) with default parameters^77^. The program used the multiple Fast Fourier Transform weighted stitching algorithm (MFWS) to splice the morphologic nucleus staining images of the tissue sections. Briefly, image splicing and registration were performed based on “track lines” (the marking lines designed on the three-dimensional Stereo-seq chip). The morphologic staining image, the corresponding spatial gene expression map, and a deep learning model (U-Net) were used to predict the mask of the cell nucleus to obtain single-cell segmentation. The algorithm (MLCG) based on cell morphology and Gaussian Mixture Model (GMM) was used to obtain the gene expression profile at the single-cell level.

### Annotation of cell types in Stereo-seq data

We used SingleR to annotate the cell types of bin35 data, employing the reference datasets provided by SingleR^16^, which contained both neural cells and immune cells.

### Annotation of brain region in Stereo-seq chips

The bin clusters were annotated based on the *in situ* expression patterns of the brain region marker genes as well as the anatomical information from the Allen Brain Atlas (https://atlas.brain-map.org/). For a cluster spanning two brain regions, it was named by the combination of the two respective brain regions, such as MB_HB. The clusters covering multiple brain regions sporadically were annotated as “UK (Unknown)”.

### Cell annotation of specific brain regions

The expression data of a specific brain region from every chip were extracted and merged using Seurat^78^. The standard analysis procedures were performed on the integrated gene count matrix, including data normalization, identification of the top 2000 highly variable genes, PCA dimension reduction, batch correction by Harmony^79^, and clustering. It is worth noting that to avoid the influence of viral genes on clustering, we excluded viral genes from the highly variable gene list. We then used the ‘FindAllMarkers’ function (log_2_fc.threshold = 0.25 and assay=”Spatial”) to find marker genes for each cluster and used these marker genes for manual cell annotation of these clusters. For secondary clustering of specific cell types of interest (such as immune cells), a similar approach was used by first subsetting the cell population of interest and then repeating the above steps.

### Determination of virus infection profile

According to the brain region annotation results, the proportion of virus-positive bins within each brain region was calculated. To filter out possible false positive bins, only bins that contained at least two JEV genes or three viral reads were deemed as JEV-positive.

### Identification of differentially expressed genes (DEGs)

After normalization, we merged all 12 samples using Seurat’s ‘mergè function and removed batch effects. Differential expression analysis was conducted to identify genes with significant expression changes across the four timepoints: 0 dpi, 3 dpi, 5 dpi, and 7 dpi. For each pairwise comparison (e.g., 3 dpi vs. 0 dpi), Seurat’s ‘FindMarkers’ function was employed using a Wilcoxon rank sum test. Genes were considered differentially expressed based on a threshold of adjusted p-value < 0.05 and a log2 fold change > |0.25|.

### Gene Ontology (GO) enrichment analysis

Lists of DEGs were used as input for the ‘clusterProfiler’ R package to conduct GO enrichment analysis^80,81^. GO terms in the categories “Biological Process”, “Molecular Function”, and “Cellular Component” were considered. The ‘enrichGÒ function was utilized to identify overrepresented GO terms in the DEG lists. The Benjamini-Hochberg procedure was applied for multiple testing corrections and the GO terms with a corrected p-value < 0.05 were considered significant. Enrichment analysis and visualization were performed with clusterProfiler^80,81^. Online website Metascape (https://metascape.org/) was also used for pathway enrichment analysis, after inputting the genes as well as choosing their matching species (*H. sapiens* or *M. musculus*), all the analysis was conducted with default parameters.

### Gene set scoring analysis

The expression scores for gene sets of specific biological pathways were calculated using the AddModuleScore function in Seurat^78^ and GSVA (Gene Set Variation Analysis)^82^. AddModuleScore calculated a score for each bin or cell by obtaining the average expression levels of selected genes and subtracting the aggregated expression of control feature sets that were randomly selected. GSVA analysis was also used to determine whether a gene set of interest was enriched in certain cell types. The scores of interested gene sets were obtained using the “gsva’’ function in the GSVA (1.46.0) package with default parameters. The hallmarks for apoptosis, necroptosis, ferroptosis, cuproptosis, pyroptosis, and autophagy were acquired from published articles^58,83–85^ (Supplementary Table 5). The other gene sets or pathways were acquired from GSEA (https://www.gsea-msigdb.org/sea/msigdb/mouse/genesets.jsp) website or GO Database. The gene sets used for AddModuleScore and GSVA were the same.

### Cell-cell communication analysis

We performed cell-cell interaction analysis using CellChat (version 2.1.0), taking into account the spatial information of bins/cells^21^. The analysis was conducted using a Mouse database^20^. We extracted the gene expression profile of each sample from the integrated dataset. The cell type information, expression matrix information, and cell position coordinates of each sample were input into CellChat. Since the diameter of bin35 is approximately 17.5 μm, the parameter “spot.diameter” was set to 17.5. For single-cell resolution data, the “spot.diameter” parameter was set to 10. Next, we utilized the “computeCommunProb” function (”type = ‘truncatedMean’”, “trim = 0.1”, “distance.use = TRUE”, and “interaction.range = 250”) to infer cell-cell interactions between cell clusters. We retained interactions detected in at least 3 bins/cells of each cell cluster. After completing the analysis for each sample, the results were merged to compare across samples, including the ligands involved in cell interactions and their associated pathways. NicheNet (version 1.1.0) was used to infer the target genes influenced by interacting cells based on prior knowledge of potential ligand-receptor and ligand-target resources ^22^.

### Identification of microglia subpopulations

We selected those bins that contain the inflammatory associated genes including *Cd14, Adgre1, Aif1, Cd68, Cd163, Cx3cr1, Fcrls, Itgam, Itgax, P2ry12, Sall1, Tmem119, Ccr2,* and *Trem2* for cluster analysis by Seurat. We further used marker genes for microglia (*C1qa, C1qb,* and *Ccl12*) to identify clusters with microglial signals and performed a secondary clustering. Finally, we adjusted the resolution to identify subtypes of *Ccl12* microglia by differential expression of *Il2b, Spp1,* and *Cxcl13*. All the above analyses were conducted in Seurat^78^.

### Gene co-expression module analysis

Gene co-expression module analysis was performed using hdWGCNA^28^ (0.2.17) on the integrated dataset combining the same brain regions from different samples. In brief, this process involved the use of the hdWGCNA function “MetacellsByGroups” to construct metacells separately for each sample and each cell groups, with each metacell encompassing 25 bins. For all cell groups, the following functions were sequentially executed using default parameters: “TestSoftPowers”, “ConstructNetwork”, “ModuleEigengenes”, and “ModuleConnectivity”. After that, the “GetModules” function was employed to extract the genes within each module, and pathway enrichment analysis was conducted on the module genes to determine their functions. Modules enriched in pathways related to learning, memory, cognition, and movement or locomotory behavior were selected, and the corresponding genes for these pathways were identified. Further examination of the expression patterns of these functional genes in different groups was performed within the Seurat object^86^.

### Transcription factor (TF) analysis

Transcription factor regulation analysis was performed using Pyscenic^26^ (version 0.12.1). In brief, the analysis involved three main steps. Firstly, grnboost2 was employed to identify co-expression networks of transcription factors and genes based on the input cell-gene expression matrix. Secondly, RcisTarget was utilized to search for significantly enriched motifs within each co-expression module, based on its internal gene-motif and motif-transcription factor databases^26^. The significantly enriched motifs are selected for each module and used to predict target genes. By integrating the TF-gene modules and the predicted target genes, a gene regulatory network module (regulons) was constructed, which included both transcription factors and target genes. Thirdly, AUCell was employed to evaluate the activity of each regulon in each cell/bin, assessing the regulatory activity of each transcription factor and its target genes in different cellular contexts^87^.

### scRNA-seq data and analysis

Single-cell data were obtained from the GEO dataset GSE237915^13^, which contains a total of 8 samples. We excluded one sample from the healthy group due to the failure in reading the matrix of the sample ‘GSM7656311’. We sequentially read the expression matrices of the remaining 7 samples using Seurat^86^ (version=4.3.0), retained the cells with mitochondrial ratio below 25% as well as gene number larger than 200, and then merged the 7 samples into an integrated Seurat object using “Merge”. The merged Seurat objects were then normalized, high-variable genes selected, and PCA downscaled according to the default parameters, and the batch effect was removed by harmony^79^ (version=1.0). Clustering was performed using ‘FindClusters’, obtaining 13 clusters at a resolution of 0.05. The ‘FindAllMarkers’ function was utilized to find the marker genes for each cluster to perform cellular annotation and a total of 11 major cell types were annotated. After normalization, the number of virus reads (labeled as “jev-p3”) in each cell ranged between 0 and 8. Cells with a normalized virus gene count >3.5 were set as virus-positive.

### Correlation assessment between scRNA-seq and Stereo-seq datasets

The microglial subsets from single-cell omics and spatial transcriptomics were analyzed separately using the “FindAllMarkers” function to identify differentially expressed genes, with an adjusted p-value < 0.05 and log fold change (logFC) > 0.25. The enrichment scores for the microglial subtypes across both datasets were calculated using the “enricher” function from the “clusterProfiler” package to assess the correlation between the two datasets^80,81^.

### Host and virus interaction analysis

We used an unsupervised sequence embedding encoding method (doc2vec) combined with a random forest classifier^29^ to predict the interactions between the 12 proteins encoded by JEV (Accession Number: L48961) and 20,433 human proteins reviewed by SwissProt. To obtain high-confidence results, the specificity parameter was set to 0.95. The database of mouse transcription factor information was collected from the SCENIC software (https://github.com/aertslab/pySCENIC/tree/master/resources), while the information of transcription factors and their target genes was obtained from TRRUST database (https://www.grnpedia.org/trrust/).

## Data availability

The Stereo-seq data supporting the findings of this study have been deposited into CNGBdb (https://db.cngb.org/cnsa/) under the accession number of STT0000076 (https://db.cngb.org/stomics/).

## Ethics statement

All animal experiments were approved by the Institutional Animal Care and Use Committee of Tsinghua University and conducted following their guidelines (approval number: 22-CG3) and Beijing Genomics Institute, Shenzhen, China (BGI-IRB A21005).

## Acknowledgement

This study was supported by grants from the National Natural Science Foundation of China (82341118, 82341082, and 32188101), the National Key Research and Development Plan of China (2021YFC2300200, 2022YFC2303200, 2021YFC2302405, and 2022YFC2303400), Shenzhen San-Ming Project for Prevention and Research on Vector-borne Diseases (SZSM202211023), Yunnan Provincial Science and Technology Project at Southwest United Graduate School (202302AO370010), the New Cornerstone Science Foundation through the New Cornerstone Investigator Program, and the Xplorer Prize from Tencent Foundation. We thank China National GeneBank for providing sequencing services for this project. We would also like to thank DCS Cloud (https://cloud.stomics.tech) for providing the computational resources and software support.

## Extended data

**Extended Data Fig. 1.**
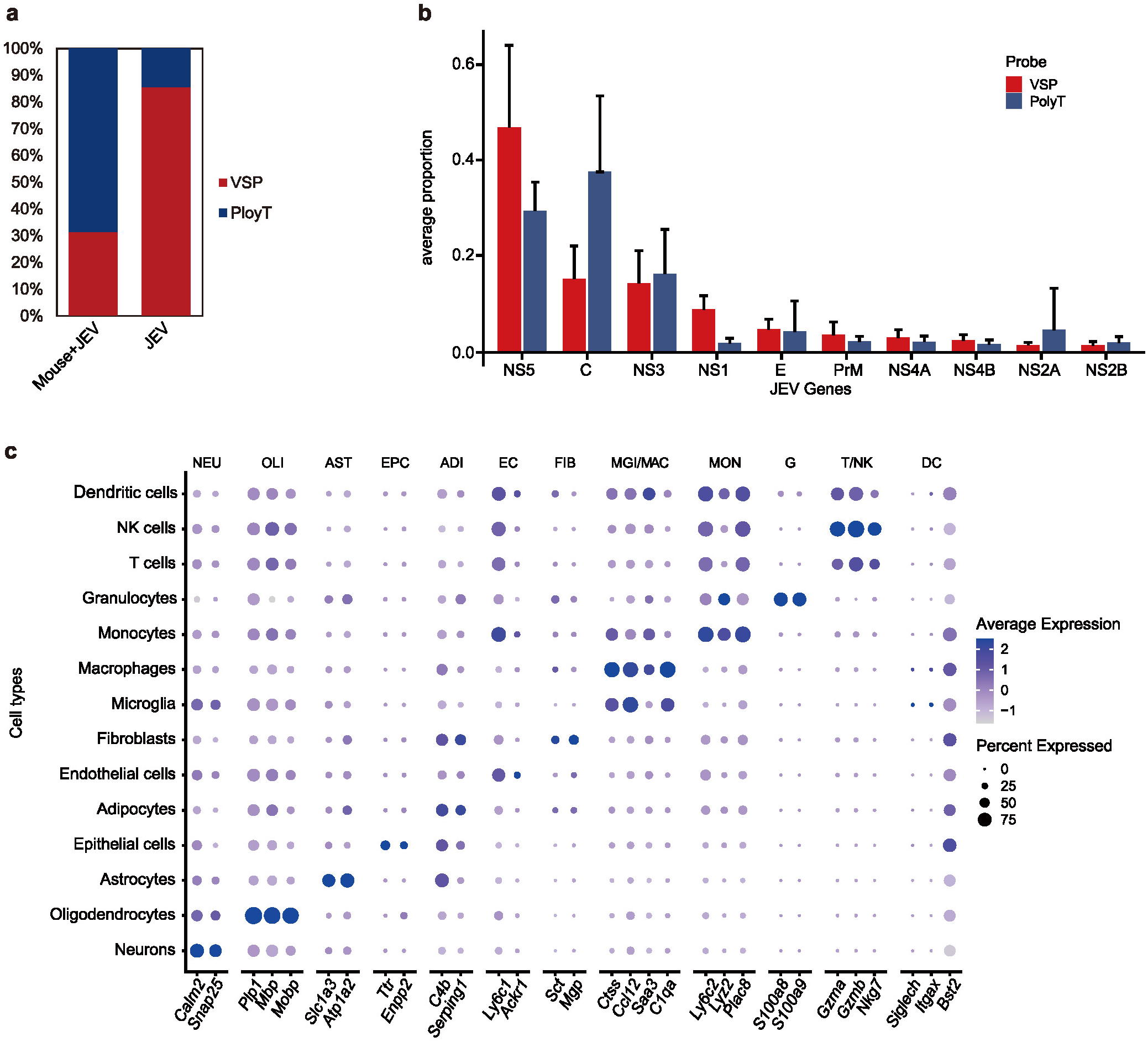
Customized Stereo-seq chips capture both JEV and mouse transcriptome. **a,** Bar plot showing the ratio of mouse and JEV transcripts captured by polyT and virus-specific probe (VSP). Both polyT and VSP can capture a low amount of non-targeted RNA through unspecific hybridization. **b,** The detection rates of each JEV coding gene by polyT and VSP. **c,** Dot plot showing the marker gene expression for each cell type in sample D7_1 (Stereo-seq bin35 data). Neu, Neurons; OLI, Oligodendrocytes; AST, Astrocytes; EPC, Epithelial cells; ADI, Adipocytes; EC, Endothelial cells; FIB, Fibroblasts; MGI/MAC, Microglia/Macrophages; MON, Monocytes; G, Granulocytes; T/NK, T cells / NK cells; DC, Dendritic cells.

**Extended Data Fig. 2.**
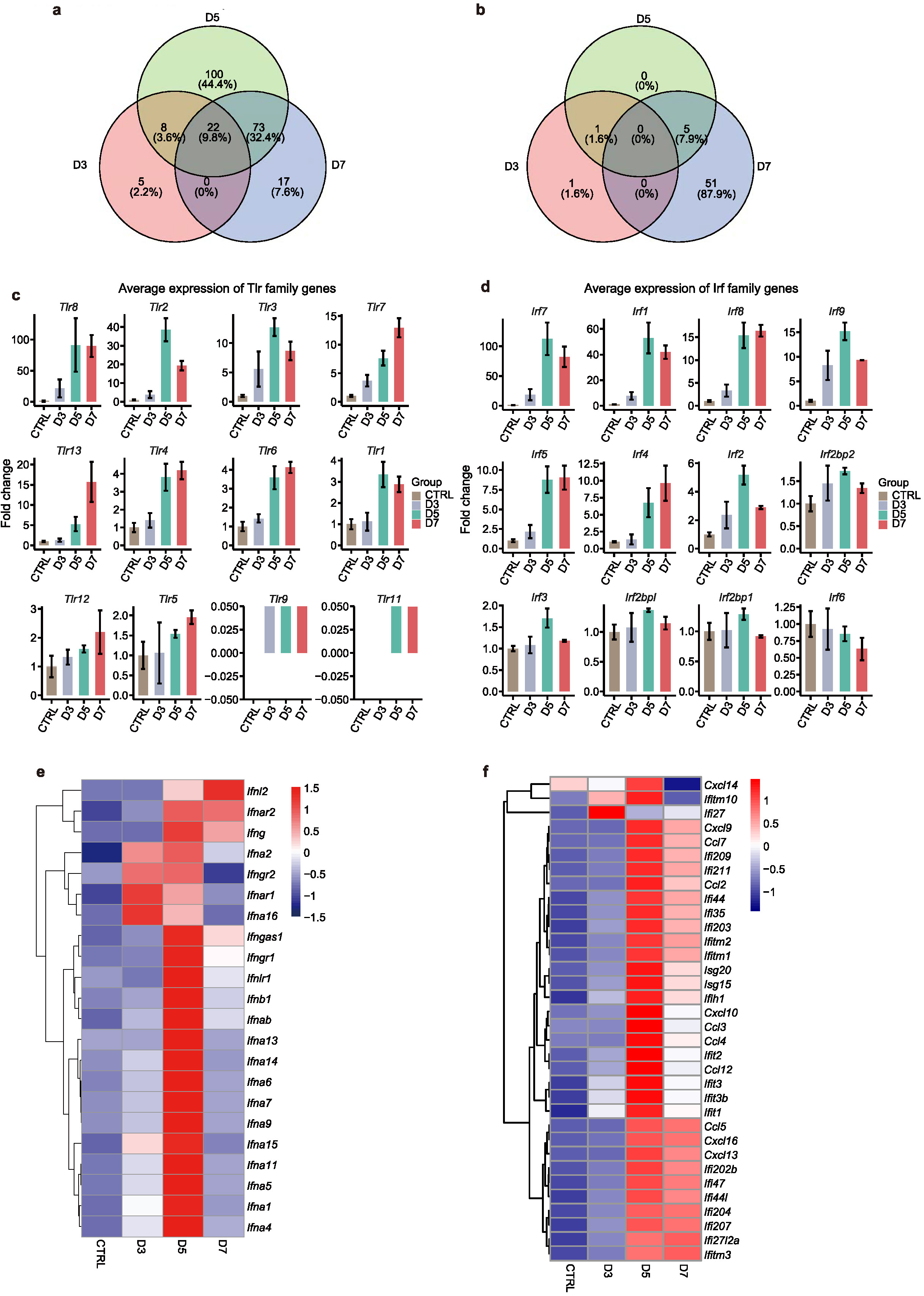
Immune response against JEV infection in the mouse brains. **a-b,** Venn plot quantifying the shared upregulated (a) and downregulated (b) DEGs in different infection groups versus control (Stereo-seq bin35 data, n=3 for each group). **c,** Bar plot showing gene expression fold changes of interferon regulatory factors (Stereo-seq bin35 data, n=3 for each group). **d,** Bar plot showing gene expression fold changes of Toll-like receptor family members (Stereo-seq bin35 data, n=3 for each group). **e,** Heatmap showing the expression of interferon gene family members in different timepoints (Stereo-seq bin35 data, n=3 for each group). **f,** Heatmap showing the gene expression of inflammatory cytokines in different timepoints (Stereo-seq bin35 data, n=3 for each group).

**Extended Data Fig. 3.**
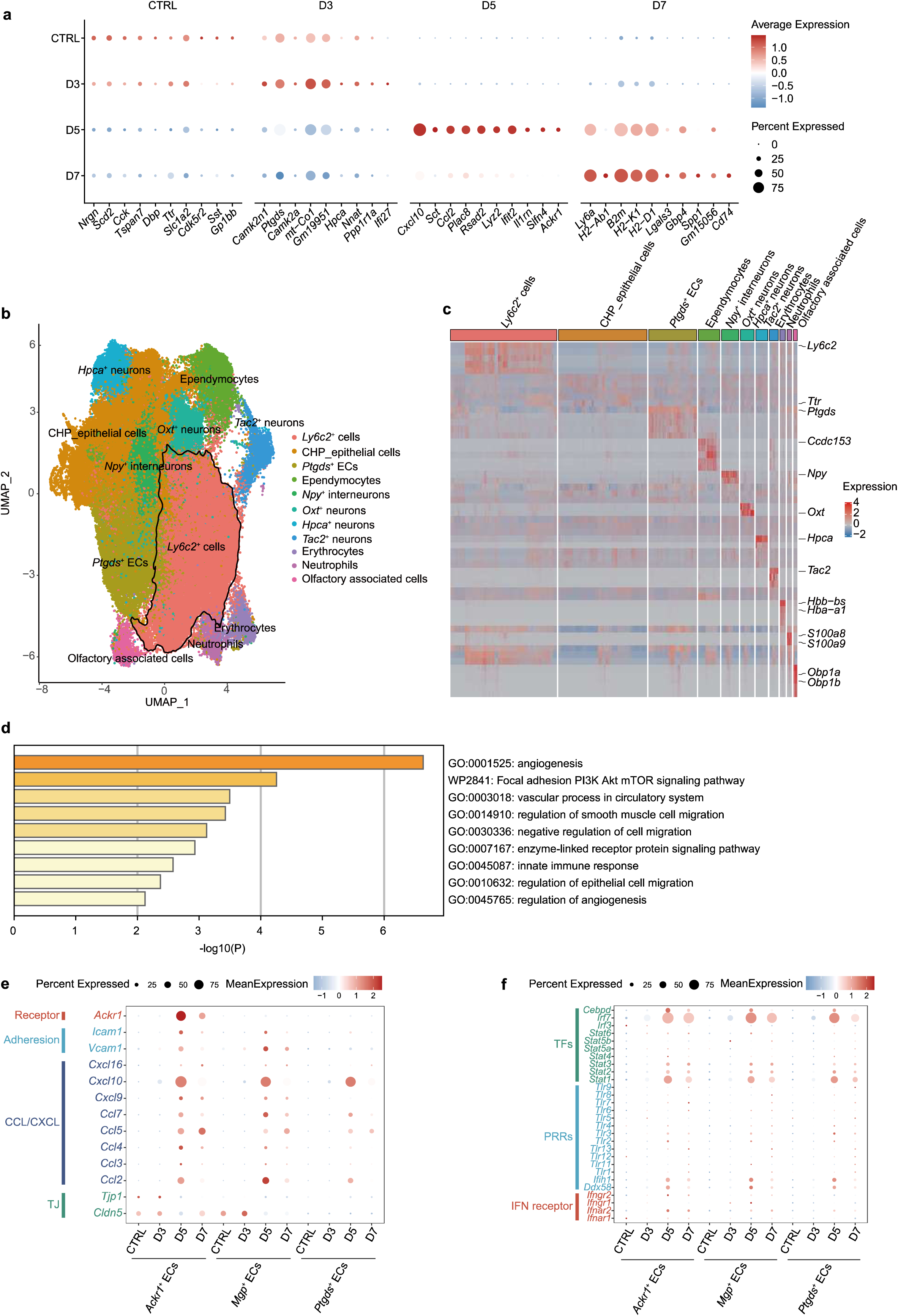
Differentially expressed genes (DEGs) and main cell type annotation in the meninges. **a,** Top10 DEGs identified in control and infection groups (Stereo-seq bin35 data of meninges, n=3 for each group). **b,** UMAP showing the major cell types identified in meninges of all samples (Stereo-seq bin35 data of meninges, n=12). *Ly6c2*^+^ cells were selected for fine annotation. **c,** Heatmap showing the marker genes for the cell types shown in (b). **d,** Enrichment analysis of the marker genes of *Ackr1*^+^ endothelial cells (ECs) identified in Fig. 3a. **e,** Bubble chart showing the gene expression for tight junction, adhesive molecules, and chemokines among different groups in different EC subtypes (Stereo-seq bin35 data of meninges, n=12). **F,** Bubble chart showing the gene expressions of transcription factors (TFs), pattern recognition receptors (PRRs), and interferon receptors in different EC subtypes.

**Extended Data Fig. 4.**
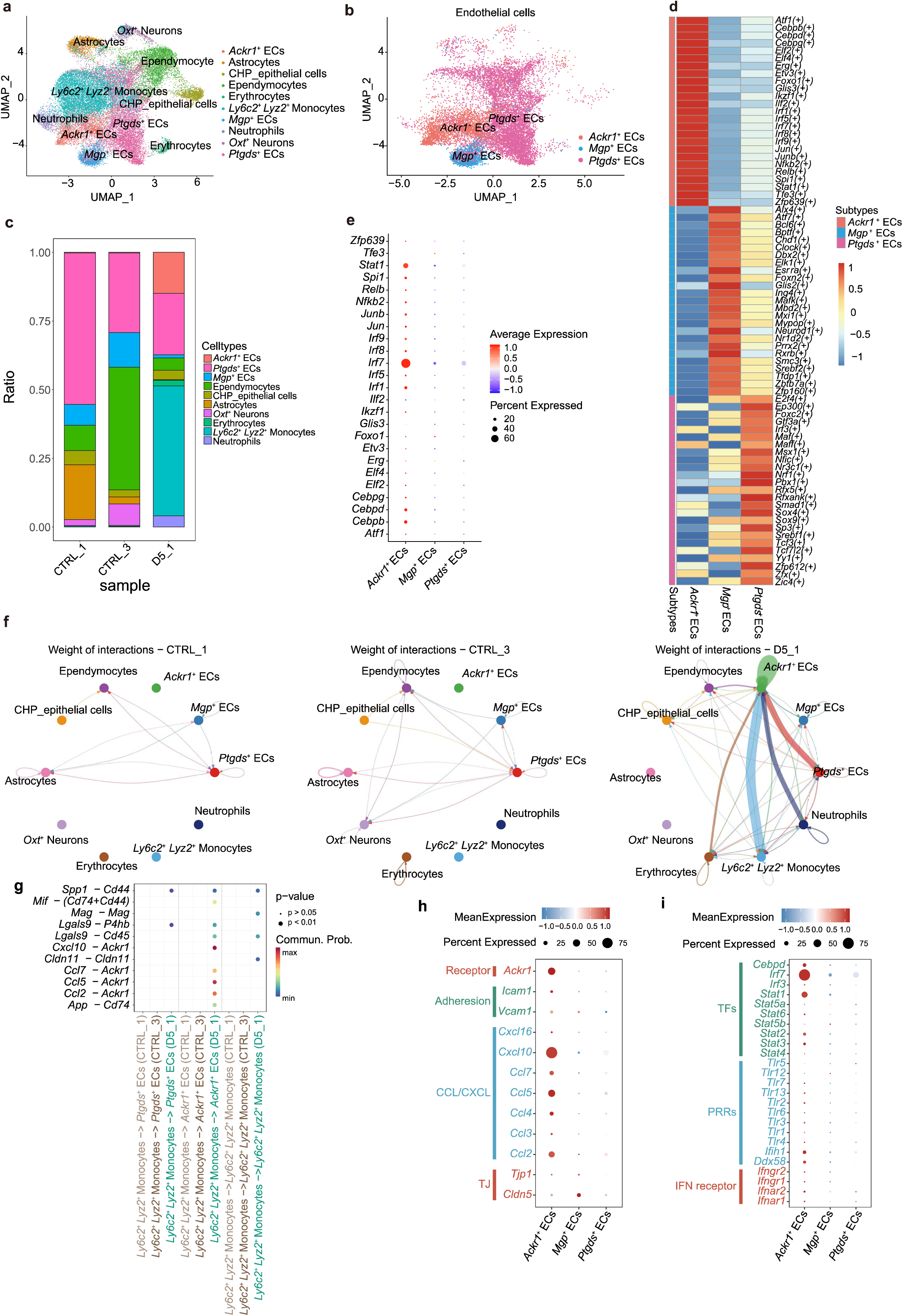
Characterization of *Ackr1^+^* ECs with Stereo-seq segmented cell data. **a,** UMAP showing the cell types in meninges of Stereo-seq segmented cell data (n=3: CTRL_1, CTRL_3, and D5_1). **b,** UMAP showing the three types of endothelial cells (ECs) in meninges of Stereo-seq segmented cell data (n=3: CTRL_1, CTRL_3, and D5_1). **c,** Heatmap showing the activity of transcriptional factors among three EC types (n=3: CTRL_1, CTRL_3, and D5_1). **d,** Dot plot showing the expression levels of the transcriptional factors specific to *Ackr1^+^* ECs (n=3: CTRL_1, CTRL_3, and D5_1). **e,** Stacked histogram of changes in cell proportions of meninges based on Stereo-seq segmented cell data (n=3: CTRL_1, CTRL_3, and D5_1). **f,** The cell-cell interactions among all cell types in meninges of sample CTRL_1, CTRL_3, and D5_1. **g,** Bubble chart showing the ligand-receptor pairs between *Ly6c2*^+^*Lyz2*^+^ monocytes and ECs (n=3: CTRL_1, CTRL_3, and D5_1).

**Extended Data Fig. 5.**
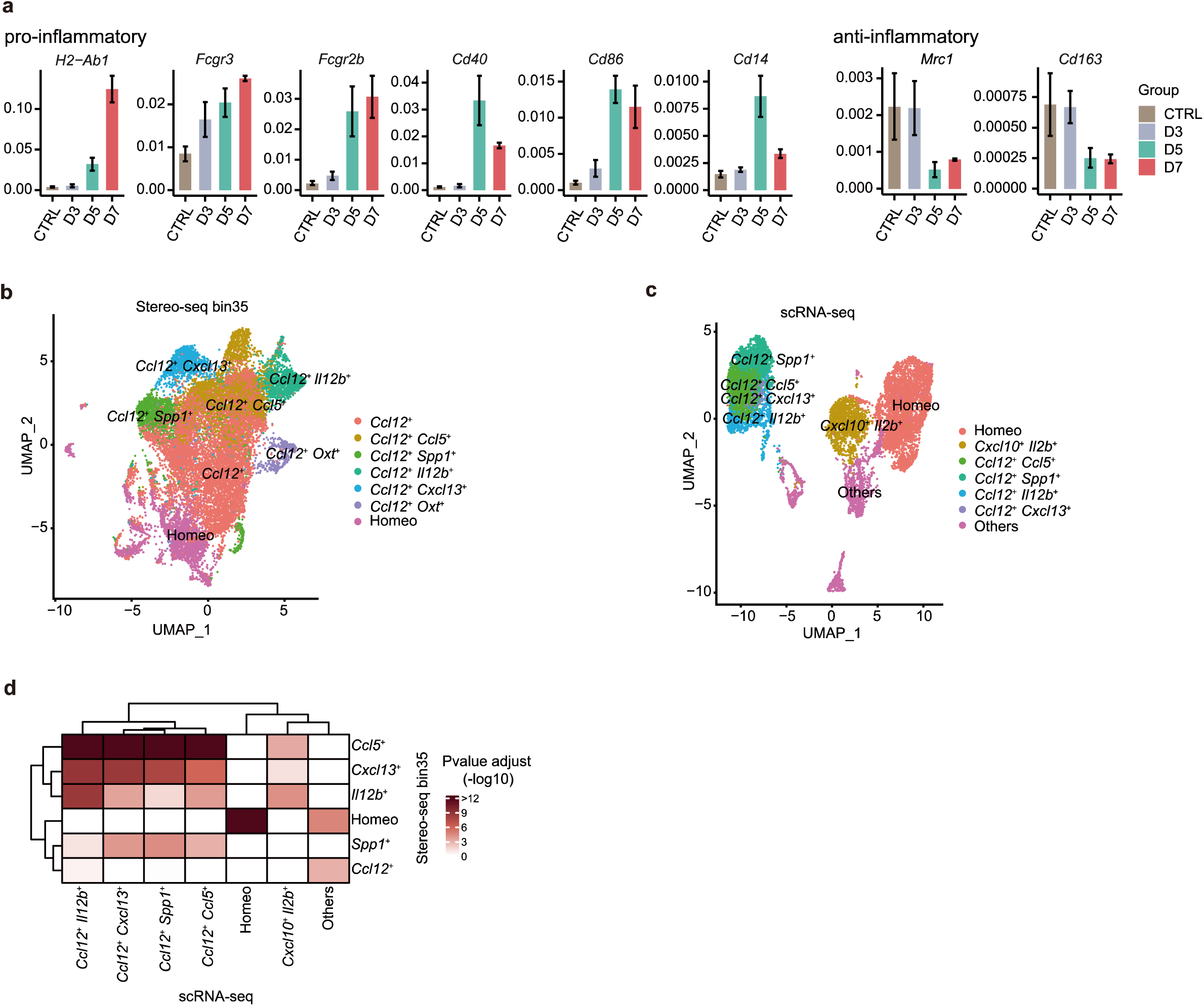
Identification of *CC12^+^* microglia. **a,** Bar plot showing the mean expression of pro-inflammatory and anti-inflammatory markers genes in microglia of the control and infection groups (Stereo-seq bin35 data, n=3 for each group). **b-c,** UMAP plots showing the microglia subpopulations in Stereo-seq bin35 data (b, n=12) and scRNA-seq data (c, GSE237915, n=8). **d,** Correlation analysis of microglia subpopulations between Stereo-seq bin35 and scRNA-seq datasets.

**Extended Data Fig. 6.**
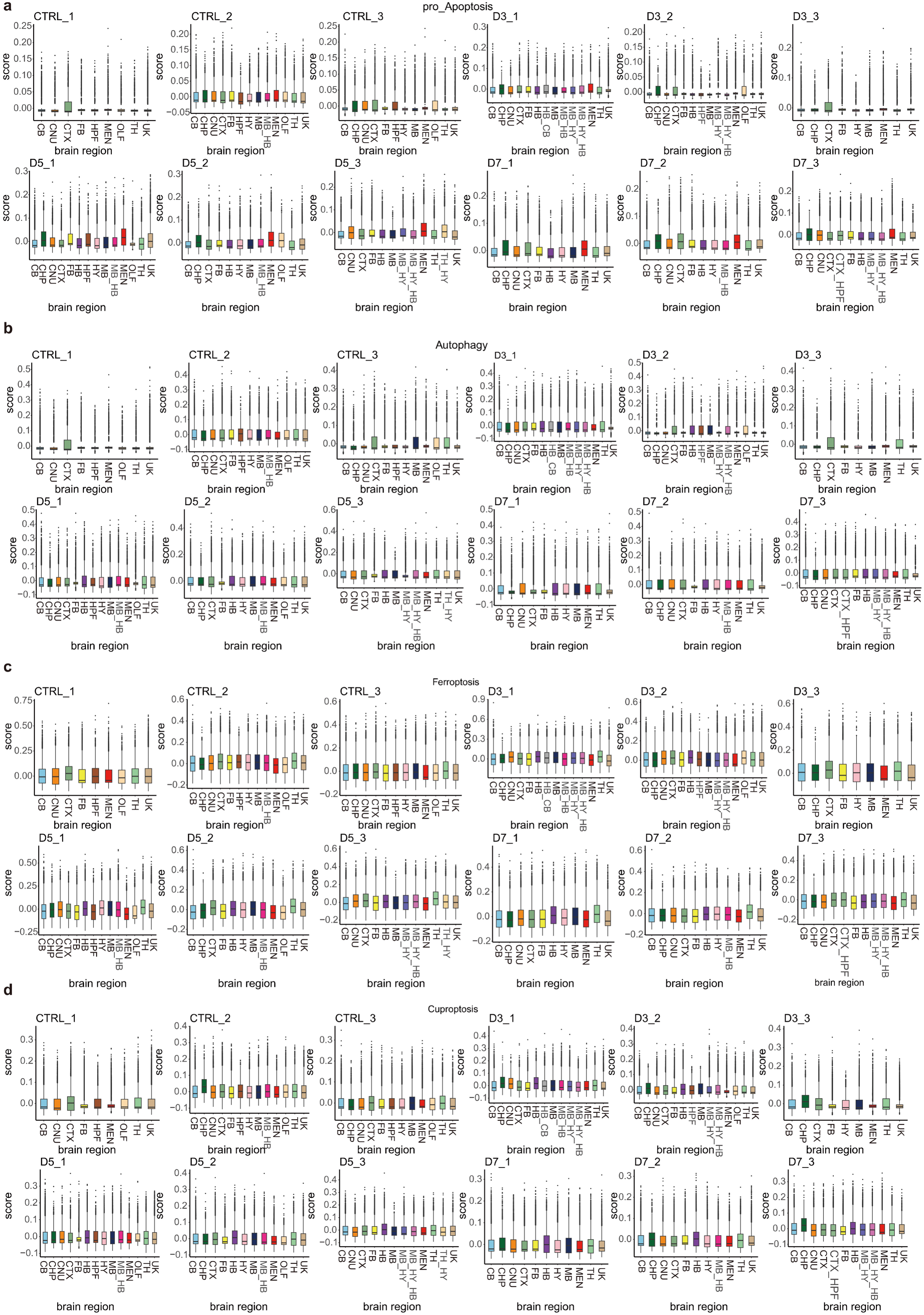
Gene signature scores of four programmed cell death pathways in mouse brains. **a-d,** The gene signature scores of pro-apoptosis (a), autophagy (b), ferroptosis (c), and cuproptosis (d) in different brain regions for each sample (Stereo-seq bin35 data, n=3 for each group).

**Extended Data Fig. 7.**
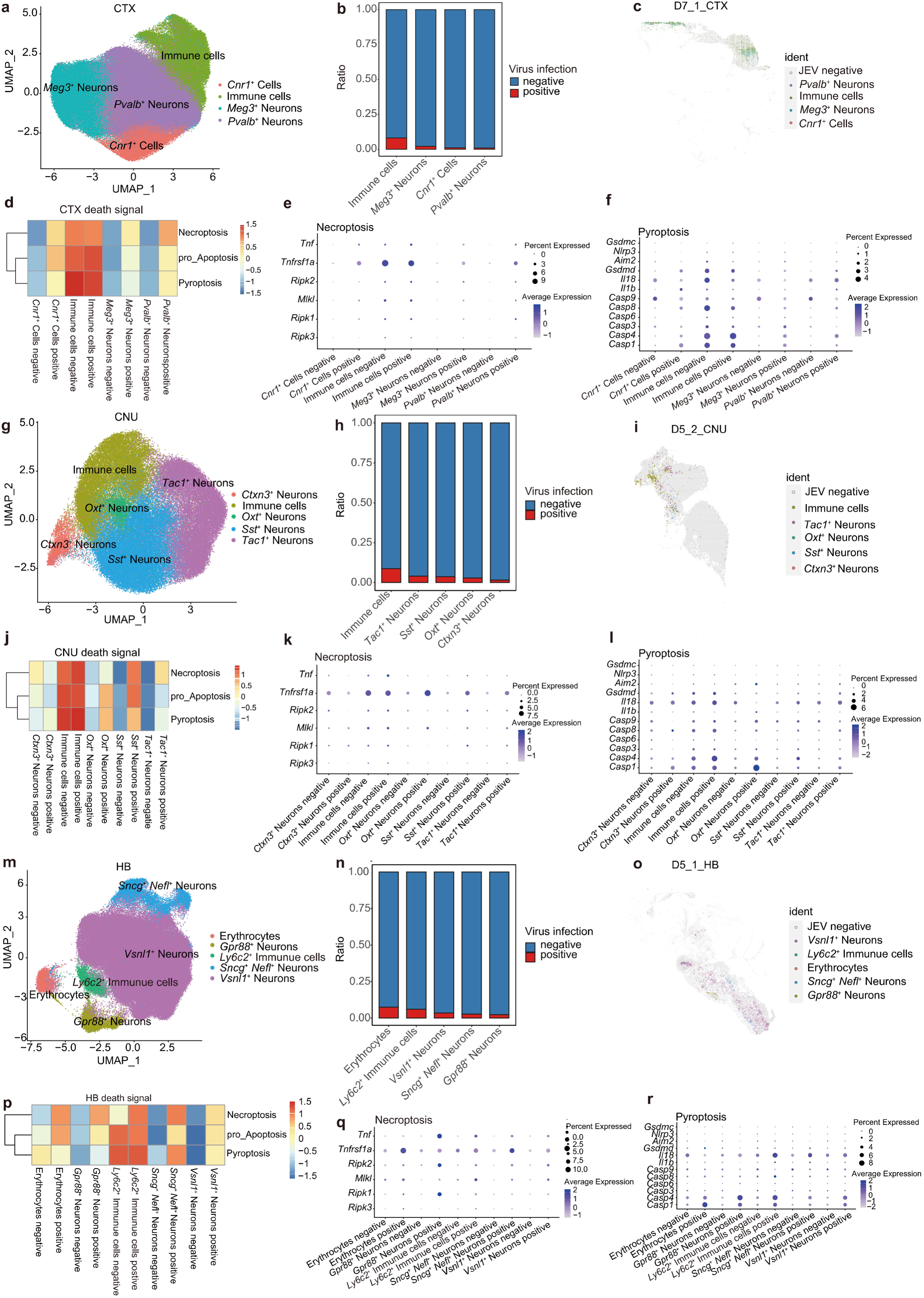
Cell death analysis in brain regions with high virus positive rates. **a,** Cell type annotation of CTX (Cerebral cortex) (Stereo-seq bin35 data of CTX, n=12). **b,** JEV positive rate among different cell types in CTX (Stereo-seq bin35 data of CTX, n=12). **c,** The spatial distribution of JEV-positive cell types in the CTX of sample D7_1 (Stereo-seq bin35 data). **d,** The GSVA scores of necroptosis, pro-apoptosis, and pyroptosis in different CTX cell types and infection states (Stereo-seq bin35 data of CTX, n=12). **e-f,** Dot plot showing the expression of essential genes involved in necroptosis (e) and pyroptosis (f) in different CTX cell types and infection states (Stereo-seq bin35 data of CTX, n=12). **g-l,** Similar analysis for cerebral nuclei (CNU) (Stereo-seq bin35 data of CNU; n=3, 2, 2, and 3 for CTRL, D3, D5, and D7 respectively). **m-r,** Similar analysis for hindbrain (HB) (Stereo-seq bin35 data of HB; n=2, 3, and 3 for D3, D5, and D7 respectively).

**Extended Data Fig. 8.**
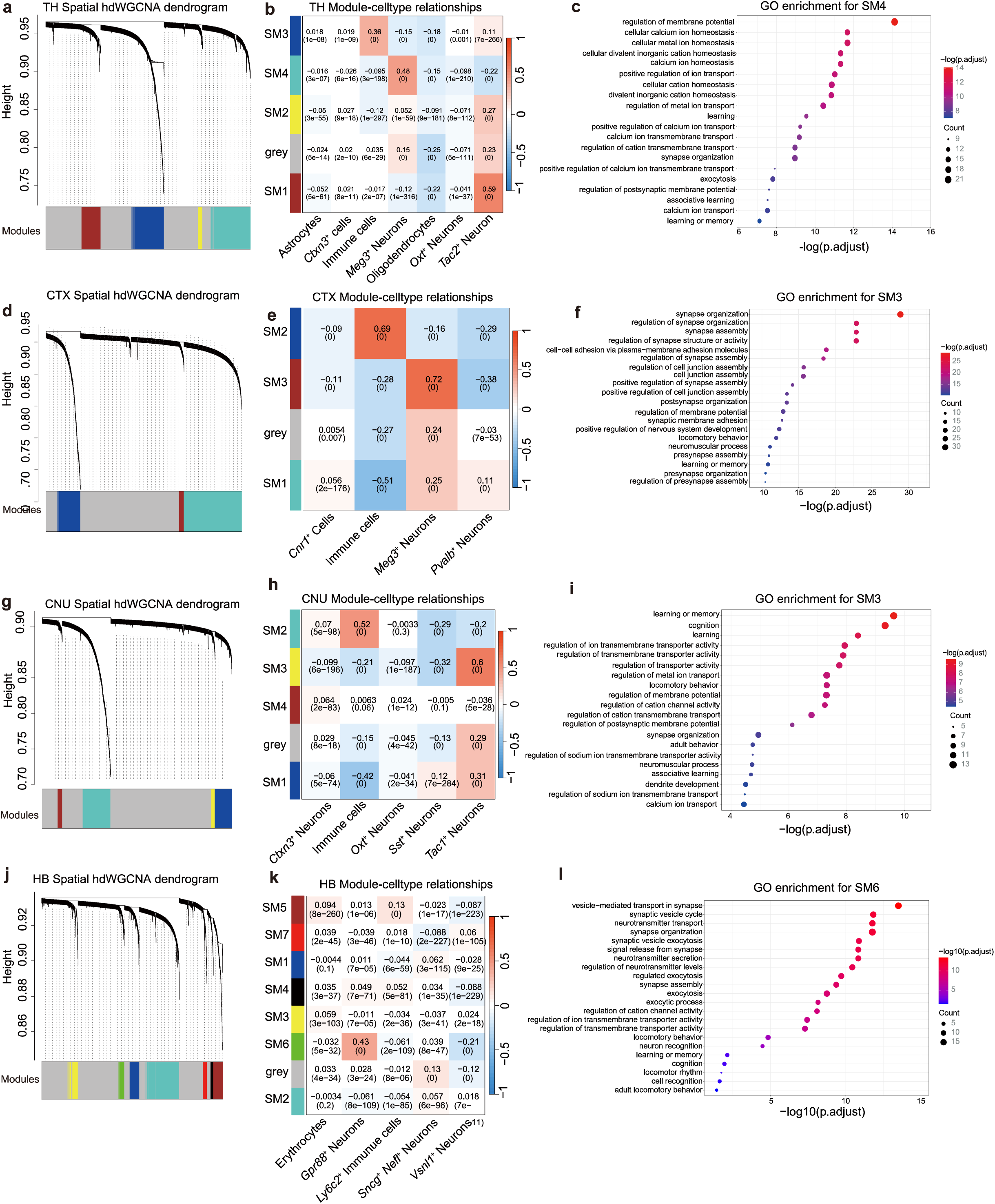
Functional module identification in brain regions with high virus positive rates. **a,** Module cluster dendrogram (top panel) and clustering of module eigengenes (bottom panel) generated by hdWGCNA based on Stereo-seq bin35 data of TH (n=12). **b,** Heatmap showing the correlations between modules and cell types in TH (Stereo-seq bin35 data, n=12). **c,** Enriched pathway in SM4 module, which is associated with *Meg3*^+^ neurons in (b). **d-f,** Similar analysis for CTX (Stereo-seq bin35 data of CTX, n=12). **g-I,** Similar analysis for CNU (Stereo-seq bin35 data of CNU, n=10, n=3, 2, 2, and 3 for CTRL, D3, D5, and D7 respectively). **j-l,** Similar analysis for HB (Stereo-seq bin35 data of HB, n=2, 3, and 3 for D3, D5, and D7 respectively).

**Extended Data Fig. 9.**
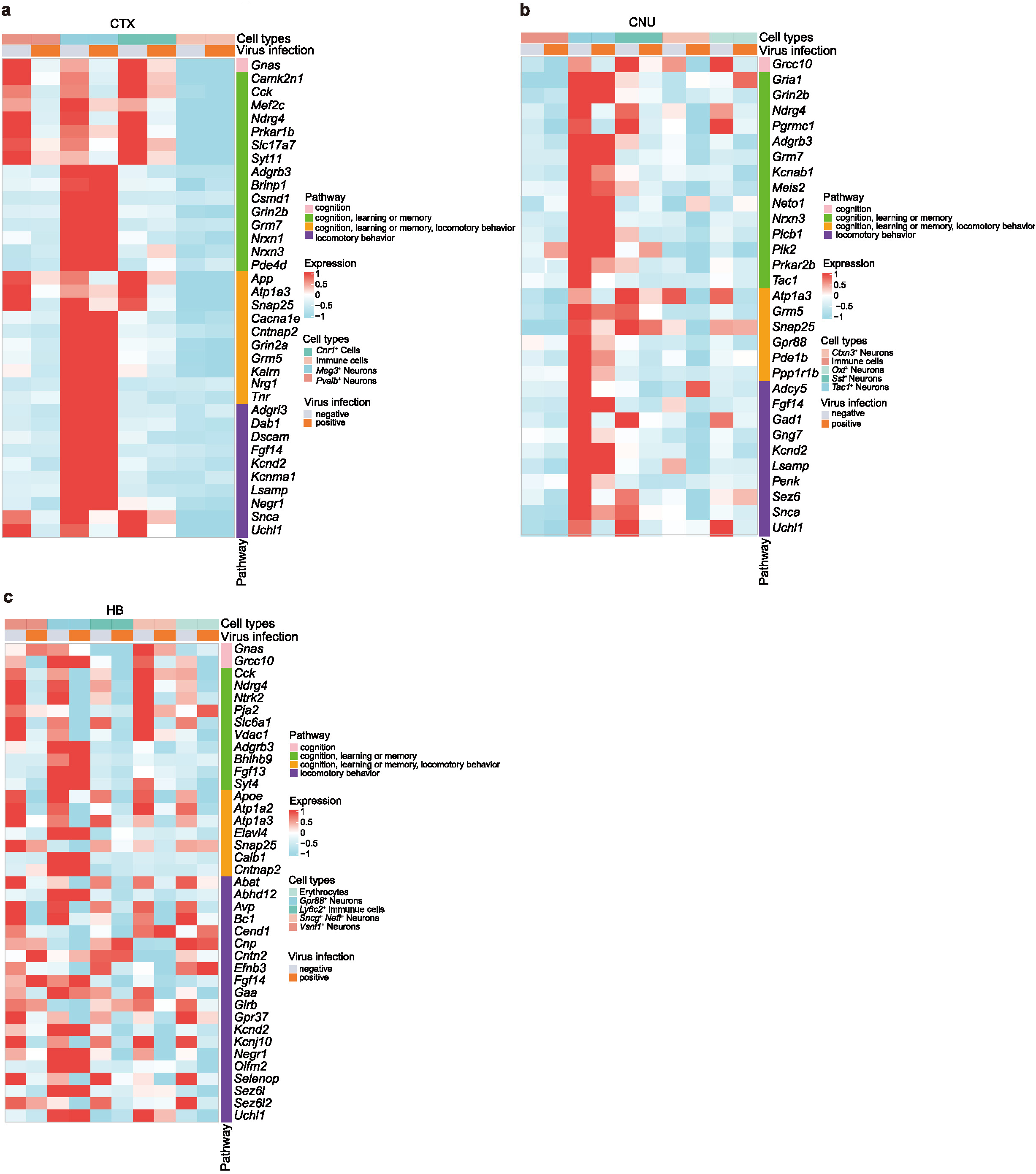
Functional gene expression in different cell types and infection states. **a-c,** Heatmap showing the expression of genes involved in cognition, learning or memory and locomotory behavior in different cell types and infection states in CTX (a), CNU (b), and HB (c) (Stereo-seq bin35 data of CTX, CNU, and HB; total n=12, 10, and 8 respectively).

**Extended Data Fig. 10.**
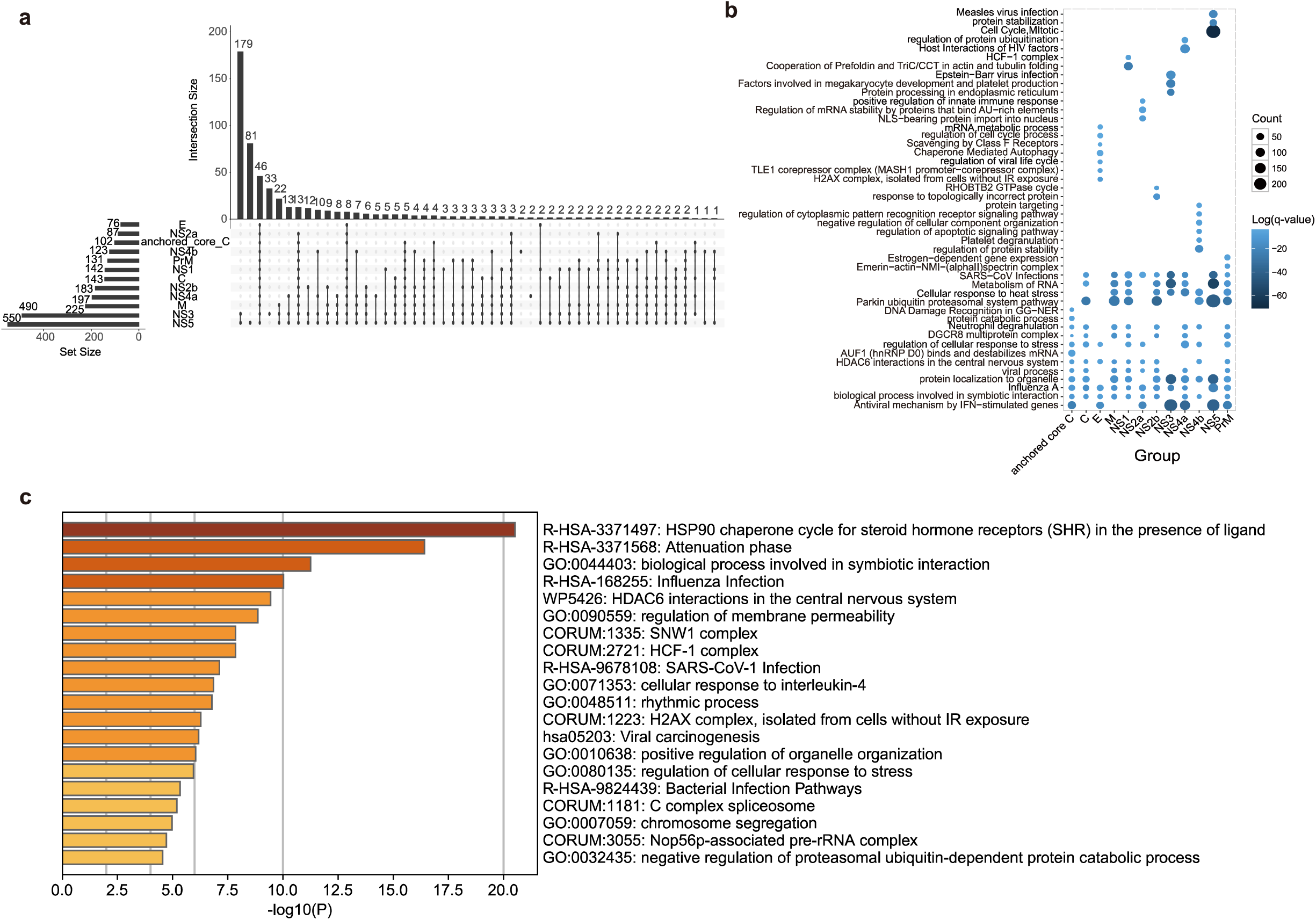
Identification of human-JEV protein-protein interactions (PPIs). **(a)** Upset plot showing the predicted number of host proteins interacting with different JEV proteins (total unique host protein=588, total viral protein=12). **b,** Dot plot showing the enriched pathways for host proteins interacting with each virus protein. Dot size shows the number of genes associated with each pathway. Dot color shows the log(q-value). **c,** Enrichment pathway of the 46 host proteins that could interact with all JEV proteins.

**Extended Data Fig. 11.**
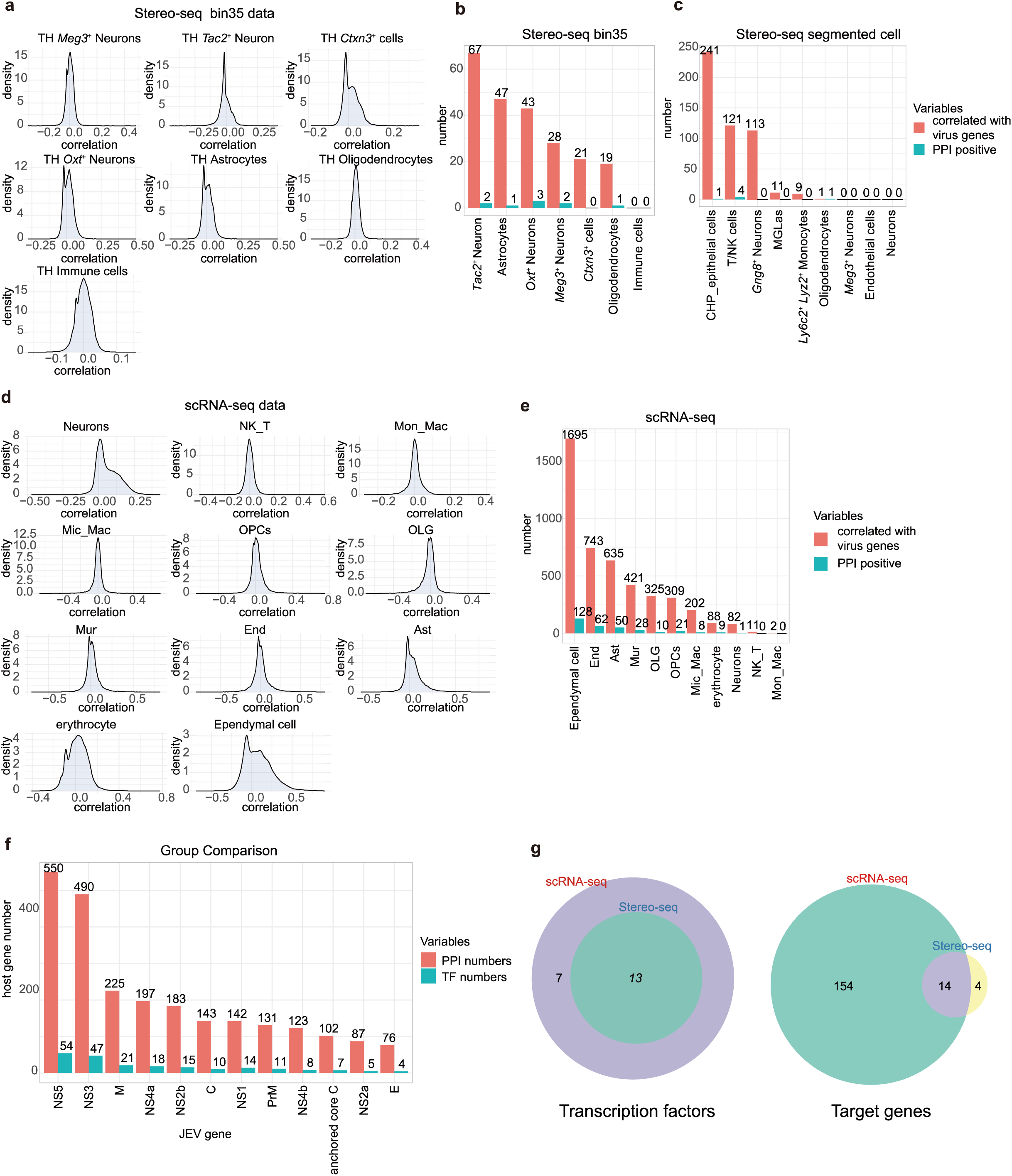
Joint analysis of PPIs with both Stereo-seq and scRNA-seq data and identification of host transcription factors associated with JEV infection. **a,** Density plot showing the correlation between virus transcripts and host genes in different cell types of TH. (Stereo-seq bin35 data, n=12). **b,** Identification of infection-associated host genes in Stereo-seq bin35 data and their relationship with host-virus PPIs. Red bar denotes the number of host genes that are correlated with virus gene expression. Blue bar denotes the number of host genes that are correlated with virus gene expression and also involved in the host-virus PPIs. **c,** Identification of infection-associated host genes in Stereo-seq segmented cell data and their relationship with host-virus PPIs (sample D5_1). **d,** Density plot showing the correlation between virus transcripts and host genes in different cell types of TH identified from scRNA-seq data (GSE237915, n=7). **e,** Identification of infection-associated host genes in scRNA-seq data and their relationship with host-virus PPIs. **f,** The numbers of human proteins and transcriptional factors that are predicted to interact with different virus proteins. **g-h,** Venn plot showing the transcriptional factors (g) and correlated target genes (h) that are involved in host-JEV interactions detected in scRNA-seq and Stereo-seq bin35 data. These intersected transcriptional factors are predicted to interact with virus protein, and the expressions of their downstream target genes were correlated with virus transcript expressions.

**Extended Data Fig. 12.**
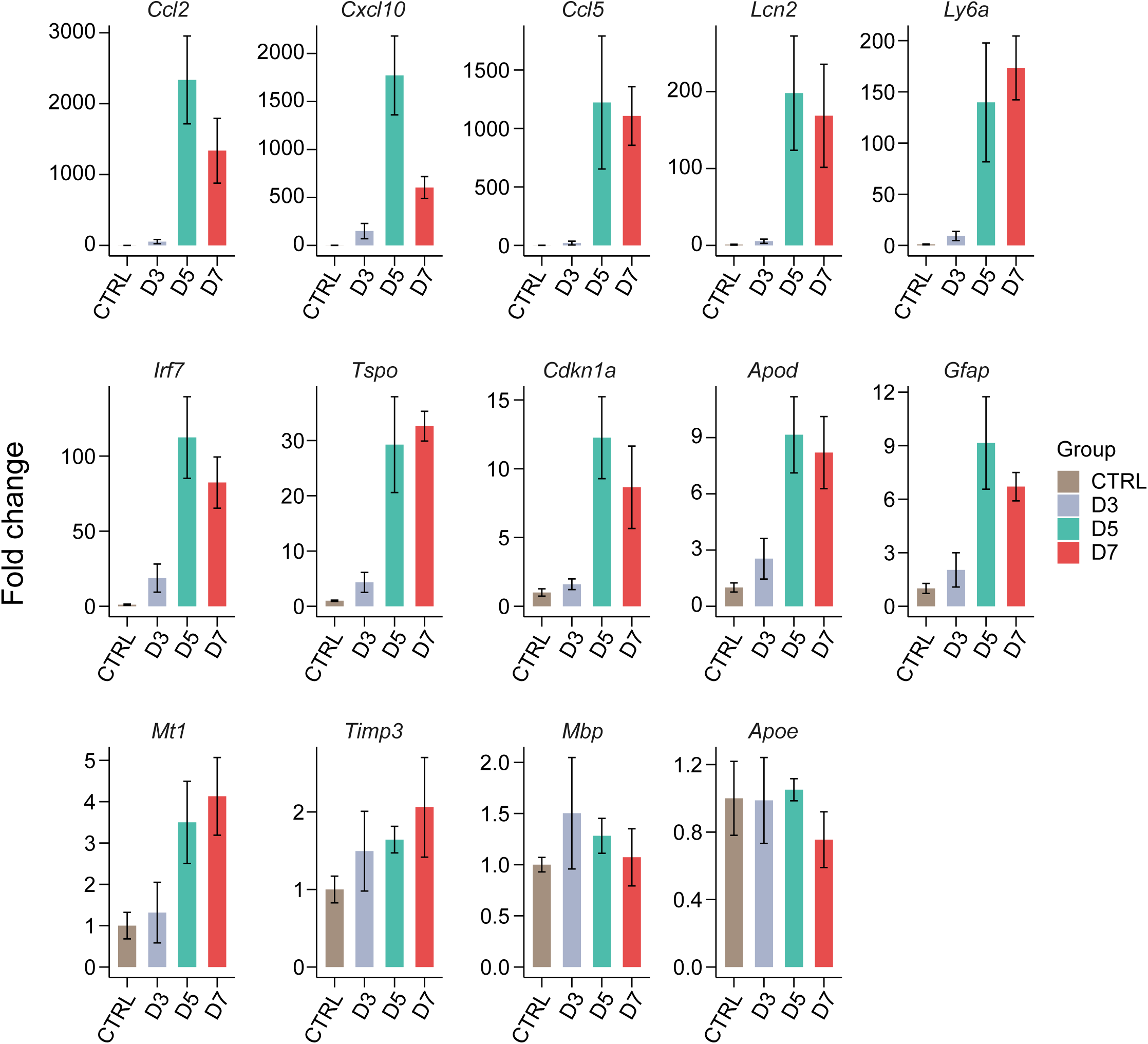
Expression of selected host genes involved in host-JEV interactions. Bar plot showing gene expression fold changes of 14 target genes in different infection timepoints compared to the control group (Stereo-seq bin35 data, n=3 for each group). The fold change calculation in the bar plot was performed by directly dividing the average gene expression of each group by that of the control group. *Mbp* and *Apoe* were negatively correlated with virus gene expression, while the other 12 genes showed positive correlations.

## Supplementary information

**Supplementary Table 1: Experimental details for the mouse samples.**

**Supplementary Table 2: Annotation markers for the mouse brain regions.**

**Supplementary Table 3: Cell types annotated in the mouse brain and their marker genes.**

**Supplementary Table 4: DEGs between control and infection groups.**

**Supplementary Table 5: Gene sets used for GSVA and gene signature score analysis.**

**Supplementary Table 6: Details of 588 host proteins predicted to interact with JEV proteins.**

**Supplementary Table 7: Enrichment pathway information for the 46 host proteins that are predicted to interact with all the virus proteins.**

**Supplementary Table 8: The host transcriptional factors that are predicted to interact with JEV proteins and their related target genes.**

